# Climate and urbanization drive mosquito preference for humans

**DOI:** 10.1101/2020.02.12.939041

**Authors:** Noah H. Rose, Massamba Sylla, Athanase Badolo, Joel Lutomiah, Diego Ayala, Ogechukwu B. Aribodor, Nnenna Ibe, Jewelna Akorli, Sampson Otoo, John-Paul Mutebi, Alexis L. Kriete, Eliza G. Ewing, Rosemary Sang, Andrea Gloria-Soria, Jeffrey R. Powell, Rachel E. Baker, Bradley J. White, Jacob E. Crawford, Carolyn S. McBride

## Abstract

The majority of mosquito-borne illness is spread by a few mosquito species that have evolved to specialize in biting humans, yet the precise causes of this behavioral shift are poorly understood. We address this gap in the arboviral vector *Aedes aegypti*. We first characterize the behaviour of mosquitoes from 27 sites scattered across the species’ ancestral range in sub-Saharan Africa, revealing previously unrecognized diversity in female preference for human *versus* animal odor. We then use modelling to show that this diversity can be almost fully predicted by two ecological factors – dry season intensity and human population density. Finally we integrate this information with whole genome sequence data from 345 individual mosquitoes to identify a single underlying ancestry component linked to human preference, with genetic changes concentrated in a few key chromosomal regions. Our findings strongly suggest that human-biting in this important disease vector originally evolved as a by-product of breeding in human-stored water in areas where doing so provided the only means to survive the long, hot dry season. Our model also predicts that changes in human population density are likely to drive future mosquito evolution. Rapid urbanization may drive a shift to human-biting in many cities across Africa by 2050.

---

Mosquitoes spread pathogens that make approximately 100 million people sick every year^1^. There are roughly 3,500 mosquito species worldwide^2^, but most cases of human disease are caused by the bites of just a few taxa that specifically target humans^3,4^. Understanding where and why mosquitoes evolve to specialize in biting humans is therefore critical for controlling and predicting disease spread.

*Aedes aegypti* provides a key opportunity to investigate this phenomenon. The globally invasive subspecies, *Ae. aegypti aegypti*, thrives in urban habitats across the American and Asian tropics, where its proclivity for biting humans makes it the primary vector of dengue, Zika, chikungunya, and yellow fever^5^. Host-seeking females take up to 95% of their blood meals from humans in nature^3^. This human-biting specialist is thought to have evolved from generalist ancestors in Africa approximately 5,000-10,000 years ago, possibly in northern Senegal or Angola^6,7^. However, in at least a few places in East Africa, the contemporary African subspecies *Ae. aegypti formosus* remains a generalist, biting a wide variety of vertebrate animals^8,9^. Little is known about the host-seeking behavior of *Ae. aegypti* in other parts of Africa, and no work to date has explicitly examined the ultimate drivers of human-biting in mosquitoes.

To address this gap, we used ovitraps to collect *Ae. aegypti* eggs from multiple outdoor sites in each of 27 locations across sub-Saharan Africa (Fig. 1a-c, Extended Data Table 1). The collections spanned a wide range of human population densities, including egg traps placed among assemblages of plastic and concrete in large cities with over 2,000 people per square kilometer, ranging to traps placed among trees and undergrowth in wild areas where mosquitoes rarely encounter human hosts (Fig. 1a,c, Extended Data Table 1). They also spanned a wide range of climates, from highly seasonal, semi-arid habitats in the northwest to forest ecosystems with year-round rain in Central Africa (Fig. 1b-c, Extended Data Table 1). We used eggs from independent traps to establish two replicate laboratory colonies for each of 23 populations, and a single colony for the remaining 4 populations (n=50 colonies total, Extended Data Table 2).

**Fig. 1.**
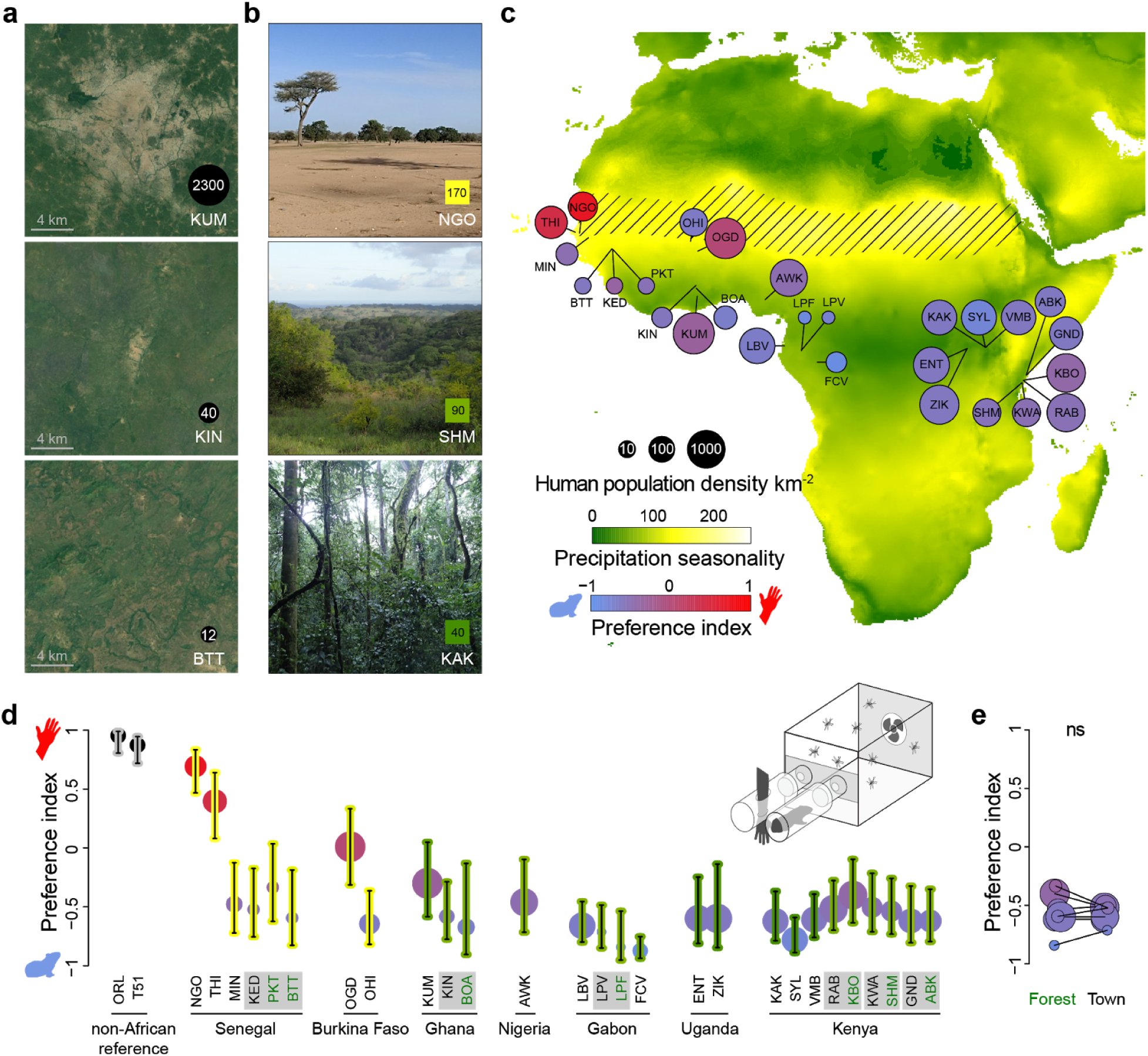
Preference for human odor varies widely in *Aedes aegypti* mosquitoes across Africa. **a-b**, Satellite images or photographs of mosquito collection localities with different human population densities (a) or levels of precipitation seasonality (b). Satellite images are from Google Earth, copyright Maxar Technologies and CNES/Airbus. **c**, Map of collection localities. Diagonal hatched lines mark the Sahel ecoclimatic zone. Extended Data Table 1 has full location names. **d**, Host preference measured in a two-port olfactometer (inset) for all African localities plus a reference colony from Thailand (T51) and a lab colony most likely to have originated in the United States (ORL) (n=3-14 trials with 25-110 females per trial; details in Extended Data Table 2). Bars indicate 95% confidence intervals. Grey boxes around location names highlight adjacent forest-town pairs (forest in green text). Circle sizes and bar colors show population density and climate, respectively, as in (c). **e**, Females from adjacent forest and town habitats did not differ in preference (*P*>0.05, ns=not significant).

## Preference for human odor varies widely across sub-Saharan Africa

Mosquitoes choose hosts based largely on body odor^4^. *Ae. aegypti* females from human-biting populations show a robust preference for human odor, while those from generalist populations often prefer the odor of non-human animals^10^. We tested the odor preference of colony females from each population in a two-port olfactometer and estimated preference using a beta-binomial mixed model that accounts for trial structure (Fig. 1d, Extended Data Fig. 1a-b, Extended Data Table 2, see Methods). The results were generalizable across different individual humans and different animal species used as stimuli (Extended Data Fig. 1c).

Most populations preferred animals, but one population from Central Africa stood out as having an extreme animal preference (Franceville, Gabon; FCV), and three from West Africa showed either no preference (Ouagadougou, Burkina Faso; OGD) or clear human preference (Thies and Ngoye, Senegal; THI, NGO) (Fig. 1d). The significance of this geographic variation (Likelihood Ratio Test *P*<2.2×10^−16^) was further supported by a strong correlation between the preference of replicate colonies from the same location (Pearson *R*^*2*^=0.60, *P*=1.5×10^−5^, Extended Data Fig. 1d). As seen in previous work, overall response rates mirrored preference, with females from animal-preferring colonies being less likely to choose either host in the assay (Extended Data Fig. 1e-f)^10,11^.

## Preference variation is explained by two ecological factors

Human-biting may be favored in areas with a high density of humans. Whether or not humans were living in the immediate vicinity of collection sites had no effect on behavior across a set of paired forest and town locations (Fig. 1e). However, linear modeling of preference variation across all locations revealed a clear effect when humans were counted within a circle of radius 20-50km (Likelihood ratio test all *P*≤0.002; compare grey and black lines in Extended Data Fig. 2a). This result supports the idea that high regional human population density helps drive mosquito preference for humans.

The population density model included latitude and longitude as covariates to control for clear geographic trends in the data (higher preference for humans in the northwest, Fig. 1c). We wondered whether climate might explain some of this additional variation. Surprisingly, when we used stepwise model selection to replace latitude and longitude with ecologically relevant climate variables (Bio1-Bio19 from the WorldClim 2 dataset; Extended Data Fig. 2b-c)^12^, the best climate variables explained more behavioral variation than human population density itself. In the final model, human population density explained 18% of variation (Fig. 2a, Likelihood Ratio Test *P*=1.0×10^−5^, density measured within 20 km radius). The strongest climate predictor was precipitation seasonality (Fig. 2b, Extended Data Fig. 2b, Likelihood Ratio Test *P*=1.2×10^−8^), a measure of how variable rainfall is from month to month. A third-degree polynomial provided the best fit (Extended Data Fig. 2a-b), helping to predict the abrupt emergence of preference for humans in the Sahel ecoclimatic zone of West Africa, where it is dry for 9 months of the year and all rainfall comes during a short, intense rainy season (Fig. 1c, 2b). A second variable, level of precipitation during the warmest quarter of the year, also contributed significantly to our model (Fig. 2c, Extended Data Fig. 2c, Likelihood Ratio Test *P*=0.014) and helped explain behavior across populations in the animal-preferring range – the bulk of our sample. Animal preference was weaker in places with less rain at the hottest time of year (Fig. 2c, Extended Data Fig. 2d).

**Fig. 2.**
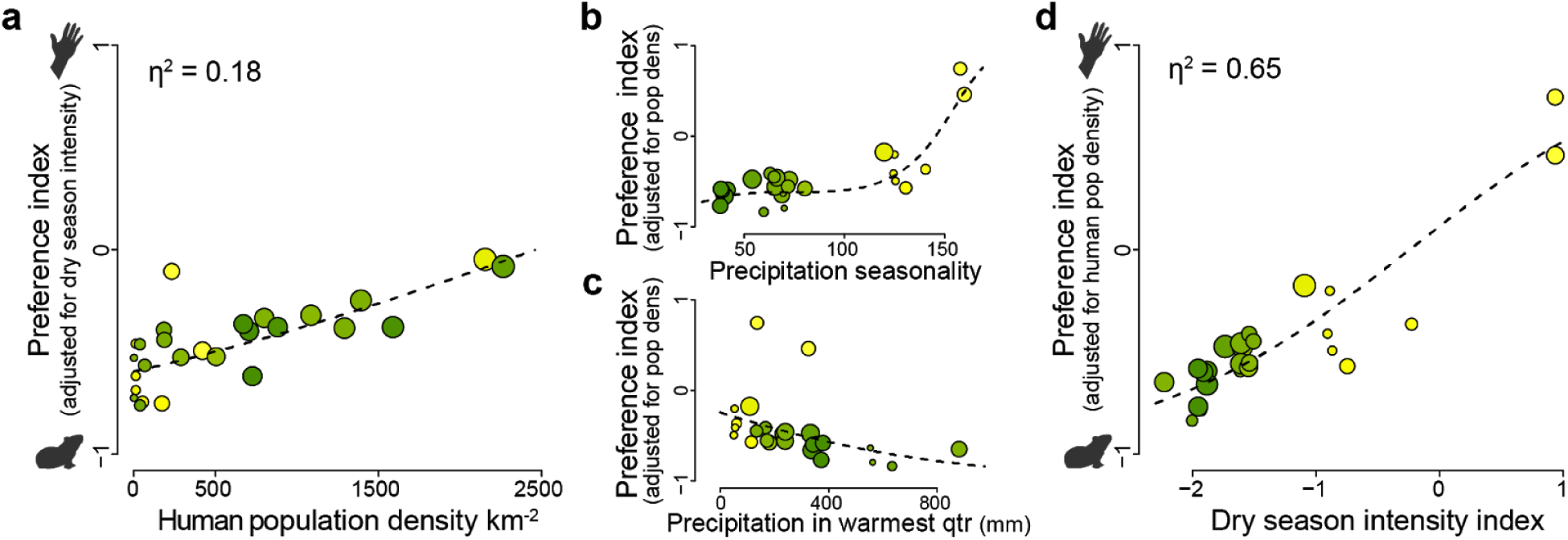
Human population density and dry season intensity explain variation in preference. **a**, Human population density explains 18% of variation in preference (linear fit, density calculated within 20km radius, Likelihood Ratio Test [LRT] *P*=1.0×10^−5^). **b-c**, Preference for humans increases in habitats with highly seasonal rainfall (b, cubic monotonic polynomial fit, LRT *P*=1.2×10^−8^) and decreases in habitats with more rain at the warmest time of year (c, linear fit, LRT *P*=0.016 for model that already includes seasonality). **d**, Overall dry season intensity (combination of variables shown in b-c) explains 65% of variation in preference (LRT *P*=3.0×10^−9^). All analyses were carried out with logit-transformed preference indices subsequently back-transformed for plotting.

Taken together, these two climate variables capture the challenges mosquitoes face during the dry season. *Ae. aegypti* lay their eggs on wet substrate just above the water line in tree holes, rock pools, or artificial containers^13^. If the eggs remain wet, they can hatch immediately. However, eggs laid in wild areas at the end of the rains must pause development and survive the duration of the dry season until rain returns – a particularly difficult challenge when the dry season is long (*i.e.* precipitation seasonality is high) and hot (*i.e.* precipitation is low at the warmest time of year)^13,14^. Human water storage helps *Ae. aegypti* in harsh environments by providing a year-round aquatic niche for larval development. We put the two climate variables together into a single index of dry season intensity that explains 65% of variation in host odor preference across Africa (Fig. 2d, Likelihood Ratio Test *P*=3.0×10^−9^). These findings point to long, hot dry seasons as a key selective factor driving *Ae. aegypti* specialization on human hosts, likely as a by-product of dependence on human-stored water for breeding^15,16^.

## Preference for humans within and outside Africa has a single genomic origin

Females of the globally invasive human specialist subspecies are characterized by light scaling on the back of the abdomen (first tergite, Fig. 3a inset, Fig. 3b grey populations)^17,18^, and previous work documented this trait in the Sahel of northern Senegal where we observed preference for humans^15,19^. We therefore wondered whether it might be linked to behavior in a continuous way across our full sample set. Indeed, abdominal scaling was strongly correlated with preference for humans (*R*^*2*^=0.81, *P*=6.7×10^−10^, Fig. 3a-b). The trend was driven not only by the most extreme variation in Senegal and outside of Africa but also by more modest variation in other regions (*R*^*2*^=0.46, *P*=0.002, Senegal and non-African reference populations excluded).

**Fig. 3.**
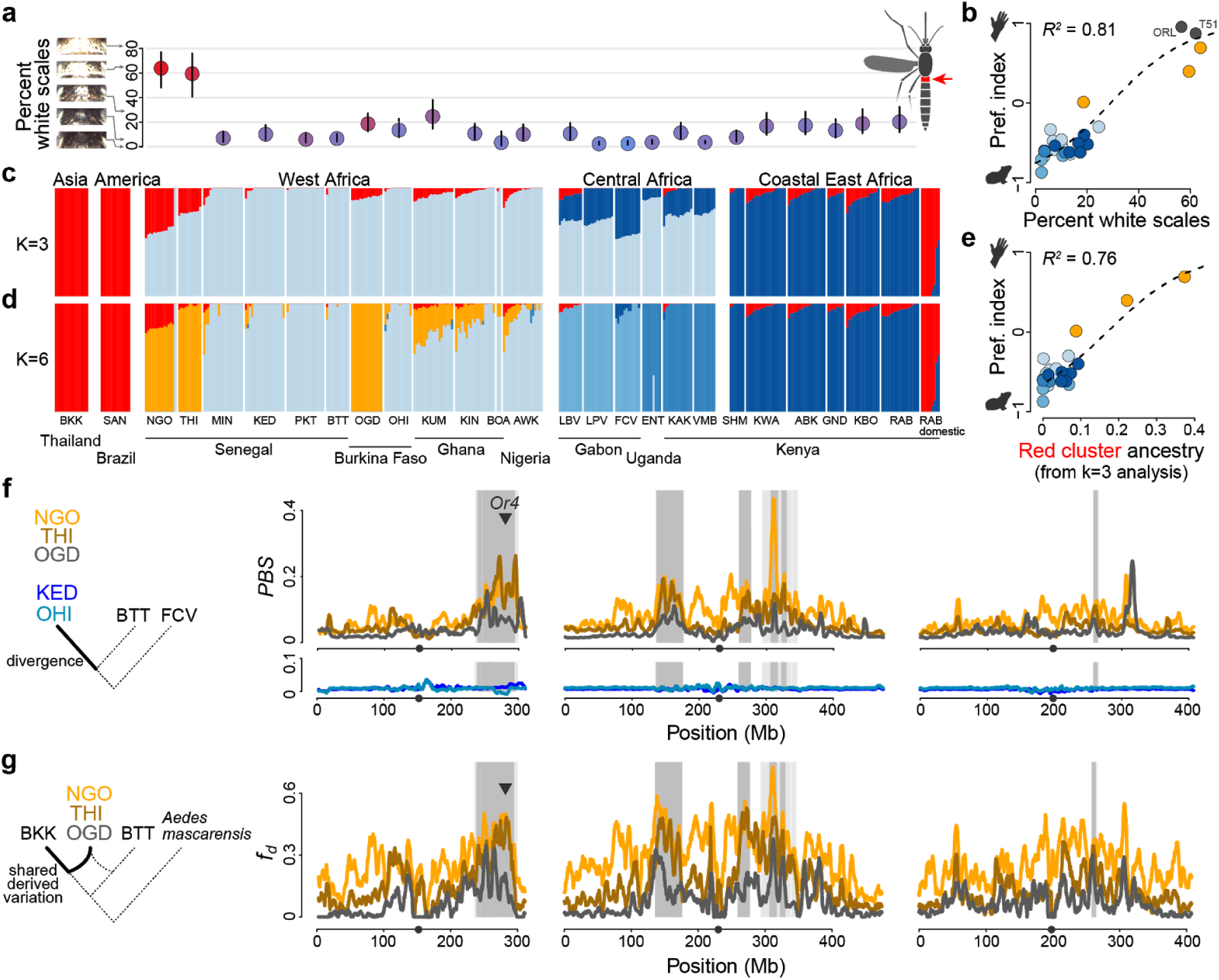
Specialization on humans has a single genomic origin and is associated with changes concentrated in a few key chromosomal regions. **a**, Variation in percentage of the first abdominal tergite (inset, right) covered in white scales. Circle colors indicate host preference as in Fig. 1. **b**, Morphology is strongly correlated with host preference. Dark grey dots represent non-African reference colonies (see also Fig. 1d). **c**-**d**, Bar plots showing proportion ancestry derived from k=3 (c) or k=6 (d) ancestry components for 375 mosquito genomes. **e**, Average proportion ancestry from the red cluster (k=3 analysis) is strongly correlated with preference for humans across populations. **f**, Polarized genomic divergence (Population Branch Statistic, *PBS*) along the lineages leading to three human-seeking populations (top; NGO, THI, OGD) and two animal-seeking populations (bottom; KED, OHI). Grey shading indicates regions that were outliers (permutation FDR<0.05) in all three human-seeking populations (dark grey) or in just the two populations showing strong preference for humans (NGO, THI; light grey). **g**, Proportion of derived variation shared by African human-seeking populations with non-African human specialists (*f*_*D*_). Grey shading as in (f). Grey triangle in (f) and (g) mark the location of *Or4*, an odorant receptor previously linked to preference for human odor^10^.

The morphological resemblance of human-preferring mosquitoes within and outside Africa suggests shared ancestry. To test this hypothesis, we sequenced the genomes of ∼15 individuals from 24 sites in our current study plus one site in South America and one in Asia (n=366 genomes after exclusion of relatives, ∼15x coverage). We also sequenced 9 previously collected individuals of the human-biting domestic form from Rabai^10^, which was not present when we carried out the fieldwork for this study in 2017. Analyses of overall population structure were consistent with earlier work^6,20^. The program ADMIXTURE^21^ revealed strong support for a model with three genomic clusters or ancestry components corresponding to coastal East Africa, Central/West Africa, and globally invasive human specialists (Fig. 3c, Extended Data Fig. 3a-b). The Rabai domestic form was the only African population to group unambiguously with non-African human specialists, consistent with its putative origin as a recent, localized reintroduction of non-African mosquitoes^20^. However, many populations across sub-Saharan Africa showed some level of ancestry from the human specialist component (red in Fig. 3c), and this signal was strongly correlated with preference for humans (Fig. 3e, *R*^*2*^=0.76, *P*=2.7×10^−8^) – supporting a single, shared origin for the behavior.

The shared ancestry of human-preferring mosquitoes within and outside African has two potential explanations – contemporary admixture due to back-to-Africa gene flow or ancestral relationships present before the species left Africa. Recent admixture has almost certainly occurred in coastal East Africa, where the reintroduced domestic form once thrived. However, a recent exome study suggested that a supposedly highly ‘admixed’ population from the Sahel region of West Africa may instead be ancestral to bottlenecked, non-African populations^7^. Consistent with this interpretation, the three most human-seeking Sahelian populations in our dataset (NGO, THI, OGD, Fig. 1c-d) formed a unified genomic cluster, distinct from both the globally invasive subspecies and nearby animal-preferring populations, in an ADMIXTURE analysis with six clusters (Fig. 3d, Extended Data Fig. 3a). Principal components analysis also shows these populations extending away from the West African cluster towards the globally invasive specialists (Extended Data Fig. 3c-d).

## Loci associated with specialization are clustered in the genome

Human-seeking mosquitoes in the Sahel have likely experienced selection for human-biting and other traits that help them live and breed in human environments – a process expected to drive and maintain divergence in parts of the genome harboring relevant loci. We used the Population Branch Statistic (*PBS*)^22^ to identify chromosomal regions with enhanced divergence along the branches leading to Ngoye (NGO), Thies (THI), and Ouagadougou (OGD) relative to nearby animal-preferring mosquitoes from Bantata (BTT) (Fig. 3f, FCV used as outgroup). Several regions stood out in all three populations (dark grey regions in Fig. 3f, permutation false discovery rate<0.05), including a large area at the distal end of the first chromosome containing an odorant receptor previously linked to preference for humans^10^. Interestingly, this same region showed signs of recent, strong selection and/or a chromosomal rearrangement in the geographic transition zone to highly seasonal Sahelian climates in Senegal (Extended Data Fig. 4).

Elevated divergence in key genomic regions might reflect the maintenance of human specialist ancestry in the face of gene flow from nearby animal-preferring mosquitoes. Using the *f*_*D*_ statistic^23^ we found that human-seeking Sahelian populations were indeed most likely to share derived alleles with non-African populations in outlier regions (Fig. 3g; Fisher’s exact test, all *P*<0.0001). Separate analyses of absolute differentiation between populations (*d*_*xy*_) and diversity within populations (*π*) were also consistent with the idea that outlier regions have been affected by strong selection during the establishment of human specialist ecology, followed by the accumulation of sequence differences via subsequent selection against gene flow (Extended Data Fig. 5)^24^.

While the most divergent chromosomal regions may harbor key loci of large effect, specialization on humans is likely to involve a diverse suite of behavioral and physiological traits with complex underlying genetics. Consistent with this hypothesis, a screen for genetic outliers more strongly associated with a human specialist ancestry component than expected under neutral evolution^25^ revealed thousands of significant single nucleotide variants scattered across the entire genome (n=16,782 SNPs at Bonferroni-adjusted *P*<0.05, Extended Data Fig. 6). However, the highest peaks fell in the same regions that stood out in the Population Branch Statistic scan.

## Rapid urbanization may drive a shift towards human-biting by 2050

Both climate and human population density are changing rapidly in Africa^26,27^. We therefore incorporated publicly available climate and human population projections into our model to explore how the behavior of African *Ae. aegypti* mosquitoes might be expected to evolve over the next 30-50 years (see Methods). Projected changes in relevant precipitation variables are modest (Extended Data Fig. 7) and unlikely to drive substantial shifts in preference (Fig. 4a, Extended Data Fig. 8). Rapid urbanization, in contrast, may trigger transitions to human-biting in many cities across the continent by 2050 (Fig. 4a, Extended Data Fig. 8), with the important caveat that the highest projected population densities are well above those used to fit our model (Fig. 4a).

**Fig. 4.**
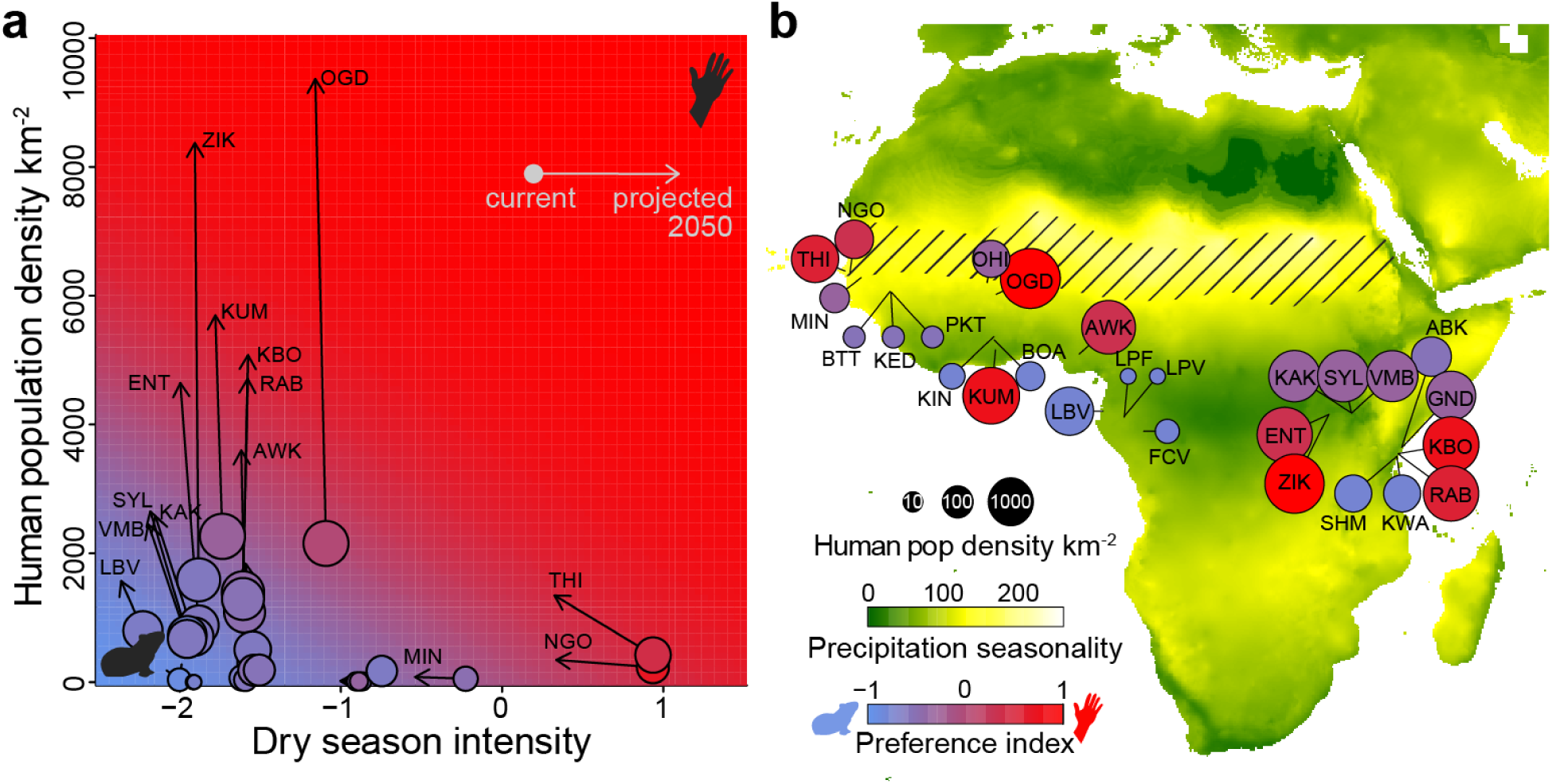
Rapid urbanization may favor a shift towards human-biting in large African cities by 2050. **a**, Current (circles) and projected (arrows) environmental parameters for each site. Circle colors indicate current host preference. Plot background color indicates preference predicted by model described in Fig. 2. **b**, Map from Fig. 1c showing projected (rather than current) human population densities (circle size), precipitation seasonality (map background color), and host odor preference (circle color) for the year 2050.

## Discussion

Our findings suggest that human-biting *Ae. aegypti* are favored in highly seasonal climates, such as the western Sahel, where laying eggs in human-stored water may be the only way to survive the dry season. Once dependent on humans for breeding sites, mosquitoes may evolve preference for humans due to trade-offs between traits that promote effective use of locally abundant human targets and those necessary for finding and biting animals^28^. There are other indications that dry season dynamics played an important role in human specialization. A decades-old study found that globally invasive specialists from Asia and the Americas, as well as a single population from the Sahel, were all more resistant to desiccation than strains from other parts of Africa^29^. Moreover, human-biting is accompanied by preference for laying eggs in human water storage vessels in at least some areas – this preference was shown to allow the introduced domestic form in Rabai, Kenya to breed throughout the dry season^30^. Beyond *Aedes* mosquitoes, dry season dynamics may have played a key role in the evolution and divergence of *Anopheles* malaria vectors in Africa^31^.

Human population density also helps predict contemporary mosquito behavior, and rapid urbanization will likely make it the most important driver of future change in Africa. However, it is unclear what will happen when human densities grow beyond the highest observed in this study, especially in the wetter and less seasonal parts of Central and East Africa. These sites may evolve strong preference for humans (Fig. 4a), as suggested by the linear trend across contemporary sites (Fig. 2a). Alternatively, the effect of human density could plateau at intermediate preference (willingness to bite humans) without driving the true specialization seen in highly seasonal environments. Despite these ambiguities, the speed and scale of ongoing urbanization argue strongly for careful monitoring of potential shifts in *Ae. aegypti* behavior (or correlated morphology/genetics) across Africa.

Our genomic data are consistent with the hypothesis that human specialization is not only favored in the seasonal Sahel, but also first arose there before seeding globally invasive populations^7^. Further work is needed to test this hypothesis – incorporating genome-wide data from a wider range of global populations. *Ae. aegypti* from Angola^6^ and northern Argentina^20^, for example, show similar patterns of ancestry to populations in the Sahel. Regardless, adaptation to humans in this mosquito is clearly associated with shifts in the frequency of a large number of variants, scattered throughout the genome but especially concentrated in a few key regions. More broadly, the tight correlations between ancestry, behavior, and environment reveal a dynamic situation playing out across the continent as a whole, with selection and gene flow fine-tuning the frequency of human-adaptive alleles, and thus levels of attraction to human hosts, according to local climate and human population density.

Our results synthesize behavioral, ecological, and genomic data to demonstrate significant, environmentally structured variation in a key disease vector. We urgently need to incorporate such variation into epidemiological models and other efforts to predict and manage the transmission of *Ae. aegypti*-borne disease in Africa.

## Methods

### Ethics and regulatory information

Mosquito eggs were collected and exported with permission from local institutions and/or governments as required (Kenya SERU No. 3433; Uganda permit 2014-12-134; Gabon AR0013/16/MESRS/CENAREST/CG/CST/CSAR and AE16008/PR/ANPN/SE/CS/AEPN) and imported to the USA under USDA permit 129920. The use of non-human animals in olfactometer trials was approved and monitored by the Princeton University Institutional Animal Care and Use Committee (protocols 1998-17 and 2113-17). The participation of humans in olfactometer trials was approved and monitored by the Princeton University Institutional Review Board (protocol 8170). All human subjects gave their informed consent to participate in work carried out at Princeton University. Human-blood feeding conducted for colony maintenance did not meet the definition of human subjects research, as determined by the Princeton University IRB (Non Human-Subjects Research Determination 6870).

### Field collections

We collected mosquito eggs in each African sampling location by distributing 20-60 ‘ovitraps’ at regular intervals across the landscape. Ovitraps consisted of 32 oz black plastic cups (The Executive Advertising), each lined with a 38 x 15 cm piece of 76 lb (34.5 kg) seed germination paper (Anchor Paper Co.) and filled with 3-8 cm of water. In all locations except coastal Kenya, the water was infused with a mixture of fresh or dry mango leaves collected from the leaf litter (n=∼20 leaves per 10 liters of water) for 1-2 days before use. In coastal Kenya and Uganda, we used tap water or made a similar infusion with twigs, bark, and leaves from unidentified broad-leafed trees. Anecdotally, water source did not appear to affect egg numbers. Each ovitrap had a hole in the side at a height of ∼3 inches to allow rainwater to drain from the trap. We attempted to spread ovitraps at approximately 100 meter intervals, but placed them at intervals as small as 10 meters in areas with limited access. We left ovitraps in the field for two nights before returning to collect egg-impregnated seed papers. We then dried the papers slowly on beds of paper towels over the course of 24 hours and stored them in airtight, whirl-pak bags during transport back to the laboratory. The only exception to this approach was at Bantata, Senegal (BTT), where we collected *Saba senegalensis* husks from the forest floor, flooded them with water, and collected hatchling larvae over the following few days. Most collections were carried out in 2017 and 2018, but Ugandan collections were carried out in 2015.

### Generation and maintenance of laboratory colonies

We hatched egg-impregnated papers from each ovitrap in separate pans of hatch broth, made by dissolving finely ground Tetramin Tropical Tablets fish food (Spectrum Brands, Inc.) in deoxygenated water (¼ tablet/liter). We continued to feed larvae Tetramin *ad libitum* through to pupation, and transferred pupae from each seed paper to separate 32 oz HDPE plastic cages (VWR). Eclosing males and females were able to mate with each other in the cages and had access to 10% sucrose solution. Other mosquito species sometimes hatched from papers alongside *Ae. aegypti* and were eventually removed from cages without hindering our breeding efforts. However, *Ae. albopictus* males are known to satyrize *Ae. aegypti* females, rendering them infertile^32^. In areas where *Ae. albopictus* was present (Nigeria and Gabon), we therefore separated the male and female pupae reared from any given seed paper and let them eclose separately before identifying adults to species and recombining *Ae. aegypti* males and females only. We set aside 2-20 adults from each population for genome sequencing, using only a single individual per ovitrap/cage where possible to reduce the probability of sequencing siblings (n=10-20 individuals for 20 locations; n=2-9 for an additional 4 locations).

We used mosquitoes from independent ovitraps to establish two replicate laboratory colonies for 23 locations and a single colony for the remaining 4 locations (Extended Data Table 2). Each colony was founded using eggs from 4-43 females, except the Lope Forest colony which was founded with eggs from a single female (Extended Data Table 2). Founding females were fed on human volunteers (see ethics subsection) and allowed to lay eggs individually on wet filter paper cones (Whatman 55 mm Grade 1 filter paper) in small shell vials (Applied Scientific Drosophila Vials, 28.5 mm diameter, 95 mm height). We gave females multiple opportunities to feed in order to ensure high feeding rates (typically >90%) and thus reduce the potential for selection on host preference. However, it was sometimes difficult to coax recalcitrant females to lay eggs in the lab. Oviposition rates ranged from 35 to 100% in the first generation. In subsequent generations, we maintained population sizes of 300-600 individuals per colony, continued to ensure blood-feeding rates >90%, and tried to maximize oviposition by forcing females into contact with wet, potting soil-infused, seed germination paper cones in small 8.5 oz HDPE plastic cups (VWR) for 2 days (30 females/cup). Eggs were dried and stored at 16°C, 80%RH for up to 6 months between generations. The only exception to these breeding procedures applied to the first 2 generations of the colony from Zika, Uganda (ZIK), which was fed on a membrane and laid eggs on cups of water placed inside a large breeding cage.

We included two reference colonies of non-African origin in behavioral and morphological studies (Extended Data Table 2). These were a colony from Thailand (T51) generated as described above and a laboratory colony originally maintained at the USDA labs in Orlando, Florida (ORL) that is of uncertain origin but was most likely supplemented for decades with local Floridian mosquitoes^33^.

### Behavior

We tested the host odor preference of 7-14 day old females that had been housed over night with access to water only (no sucrose). Different colonies were hatched on the same day, females mated freely with males after eclosion, and females were matched for age on testing days. We used a two-port olfactometer as previously described^10,11^ (Fig. 1d inset), with small modifications. Instead of using a large box fan to pull air through the device from the back of the olfactometer, we used a smaller fan to pull air through an 10.2 x 10.2 cm opening in the back panel. Instead of pulling air from the room, carbon-filtered, conditioned air was supplied to the two olfactometer ports from an independent building source. Inflow and outflow was balanced to achieve a rate of approximately 0.3m/s as measured at the traps. In each trial, 25-110 females were allowed to acclimate for 5 minutes in the large holding chamber before turning on the fan and opening a sliding door to expose them to streams of air coming from two alternative cylindrical traps and host chambers. One host chamber contained an awake guinea pig (*Cavia porcellus*, pigmented breed) or button quail (*Coturnix coturnix*). The other contained a section of the arm of a human volunteer (middle of forearm to middle of upper arm; silicone sheeting used to seal the holes through which the arm was inserted). The breath of the animal mixed with its odor in the animal odor stream. To add human breath on the human side, we asked the human subject to breathe gently through a nasal mask into the host chamber every 30 seconds. Trials lasted 10 minutes, and mosquitoes choosing to fly upwind towards either host odor stream during this time were trapped in small ports and counted at the end.

We carried out host preference trials in two main waves, with a 28-year old, European-American male serving as the human subject and one of two female guinea pigs serving as the animal subject. In the first wave, we tested second-generation colonies from Kenya and Gabon. In the second wave, we tested second generation colonies from Nigeria, Ghana, Burkina Faso, and Senegal, eighth- or ninth-generation colonies from Uganda, and two reference colonies of non-African origin. We also repeat-tested a representative set of first wave Kenyan and Gabonese colonies (RAB, VMB, FCV, LBV; by then in their fourth generation) in the second wave to ensure that results were comparable between waves. At least one of two colonies from every population included in a given wave was tested on every experimental day in order to balance random day-to-day variation with the population effects we were trying to estimate (Extended Data Fig. 1a-b). Overall, we carried out 3-4 trials for each colony, except one of two replicate colonies from BTT, KED, and KUM, which were tested only twice. This resulted in a total of 7 trials for most populations, 3-6 trials for the six populations represented by only a single colony or for which one of two colonies had fewer trials, and 14 trials for the four populations tested in both waves. In total, we carried out 206 trials including 17,856 female mosquitoes, of which 7,385 responded to one or the other host odor.

After the two main waves, we carried out a smaller set of trials with one colony from each of four representative African populations (FCV, OGD, AWK, NGO) and a wider array of host comparisons. In one set of trials, we substituted a 22-year old Nigerian-American female for the original 28-year old European-American male, and in another set we substituted a button quail for the guinea pig (Extended Data Fig. 1c, n=3-5 trials per colony x host combination).

We used a beta-binomial mixed generalized linear model as implemented in the R^34^ package *glmmTMB*^35^ to model the probability of choosing a human versus animal host for each population. This model assumes independence of individual females within trials but accounts for trial structure and the fact that preference varies more from trial to trial than is expected for a binomial model (is overdispersed) due to random sources variation (e.g. exact starting position of females within the acclimation chamber at the start of a trial, small differences in airflow between right and left ports, uncontrollable trial-to-trial variation in live host stimuli etc.). Replicate colonies and trial day were included as random factors, while population was modelled as a fixed factor. We switched the guinea pig used and the side of the human versus animal host between days such that these effects would be subsumed under the trial day random factor. We used the R package *emmeans*^36^ to extract from our glm the fitted probability of choosing a human host with 95% confidence intervals. For purposes of data visualization, we transformed each probability (p) into a preference index (PI) ranging from −1 to 1 using the formula PI=2p-1. An index of zero means the mosquitoes were equally likely to choose either host (no preference), while an index above or below zero means the mosquitoes were more likely to choose the human or animal, respectively. We used a likelihood ratio test to compare our glm to a null model accounting for day-to-day variation but not population of origin. The same beta-binomial mixed generalized linear model was used to model the probability of responding to either host (overall response rates, Extended Data Fig. 1f).

### Ecological modeling

We first compared the behavior of mosquitoes from paired forest and town populations within 5-60km of each other. In one case, a single town population (KED) was paired with two forest sites (BTT and PKT). We estimated the effect of forest habitat using a linear model that estimated preference for each pair (or group for KED, BTT, and PKT) and a coefficient for forest or town habitat. This is conceptually very similar to carrying out a paired t-test, except it allowed us to take into account the two different forest sites near KED.

We next explored the ecological factors associated with preference for humans across all populations in the sample set, again using a linear modelling framework. In this set of analyses, each population was represented by a single logit-transformed preference probability (generated by the beta-binomial model described in the previous section) and a single estimate of each ecological descriptor extracted from public datasets using the mean latitude and longitude of the ovitraps that contributed to the corresponding colony (or the mean of the two independent colony means for populations with two colonies).

While immediate habitat had no effect on behavior, we hypothesized that human population density might be relevant when calculated across a larger spatial scale. We therefore used a 2.5-minute resolution population density raster from the United Nations World Population Prospects (UNWPP, 2015 population densities adjusted to country totals)^37^ to compare the effect of density across buffers of the following radiuses: 5, 10, 15, 20, 25, 30, 35, 40, 45, 50, 60, 70, 80, 90, 100, 200, and 300 km. In a simple human population density model that also takes regional variation into account (logit(prob) ∼ human_pop_density + Latitude*Longitude), human population density had a significant effect across a wide range of spatial scales, but was strongest with ∼20-50km buffers (black line in Extended Data Fig. 2a).

To better understand the regional drivers of variation in preference, we used the WorldClim 2 bioclimatic variables (Bio1-19) as a set of candidate predictors. Because some of these variables are correlated, we also considered the predictive value of the first three principal components from a PCA analysis of Bio1-19 variation across our populations. In preliminary tests, Bio15 (precipitation seasonality) clearly showed the strongest single-variable association with preference. This was true both in a comparison of simple correlations between each variable and preference (Bio15 r=0.65) and when we included each variable in a linear model with human population density (20km buffer) (red circles in Extended Data Fig. 2b). However, the relationship with Bio15 appeared to be strongly nonlinear, if still monotonic (Fig. 2b). We therefore used a two-step procedure to model the nonlinearity. We first used the R package MonoPoly^37^ to fit monotonic polynomials of different degrees to logit-transformed preference probabilities, and then included the fitted values as an offset (i.e. removed their effects before fitting) in the linear model (logit(prob) ∼ human_pop_density_20km + offset(fitted(monotonic polynomial Bio15))). We found that a third-order monotonic polynomial significantly improved model performance, minimizing the Akaike Information Criterion (AIC). Rechecking the performance of different human population density buffers in this new model context showed that 20km yielded a much lower AIC than other buffers (light blue line in Extended Data Fig. 2a). Note that this buffer most likely reflects the balance between selection and dispersal, and is not a direct reflection of adult dispersal patterns *per se*.

We were concerned that nonlinear relationships could have obscured another better predictor in our initial survey of single-variable correlations. However, after fitting monotonic polynomials of degree 1-4 for all 19 bioclim variables and the first three PC axes, a third-order monotonic polynomial fit for Bio15 still had the lowest AIC (Extended Data Fig. 2b).

To check whether additional climate variables could further improve our model. We regressed logit-transformed preference and the other bioclimate variables on our third-order fit for Bio15 and tested if residual variation in preference could be explained by the other variables. We again used a two-step procedure to model these effects. We used the R package MonoPoly to fit monotonic polynomials of different degrees to logit-transformed preference probability residuals, and then included the fitted values as an offset (i.e. removed their effects before fitting) in the linear model (logit(prob) ∼ human_pop_density_20km + offset(fitted(monotonic polynomial Bio15)) + offset(fitted(monotonic polynomial BioX))). We found that including a linear Bio18 (precipitation in the warmest quarter) term further reduced AIC (Extended Data Fig. 2c). Because Bio18 and our fitted Bio15 polynomial relationship were modestly correlated (r=-.29) and we had selected the variables in a stepwise way, we wondered if including them in a single linear model would change our estimates of their effects. In this full, final model (logit(prob) ∼ human_pop_density_20km + fitted(monotonic polynomial Bio15) + Bio18), the estimated coefficient for the fitted Bio15 component was close to 1 (1.04), indicating that fitting the effects sequentially or together didn’t make a major difference, but we used the model where both were fit together going forward. Using both Bio15 and Bio18 as covariates, we again found that using a buffer of 20km for calculating population density yielded a much lower AIC than other buffers (Extended Data Fig. 2a).

Bio15 and Bio18 both have clear connections to the length and temperature of the dry season - an important factor in survival of dormant *Aedes aegypti* eggs. We therefore combined them into a single Dry Season Intensity Index by adding together the fitted Bio15 and Bio18 terms for each location. This simple transformation yields a single linear climate term that predicts the host odor preference of mosquitoes from a given location.

### Morphological analyses

We pinned 7-29 female mosquitoes from each location (field-collected [AWK, BOA, FCV, KED, KIN, KUM, LBV, LPV, MIN, NGO, OGD, OHI, PKT, THI] or lab colony [ABK, BTT, ENT, GND, KAK, KBO, KWA, LPF, ORL, RAB, SHM, T51, VMB, ZIK]) as previously described (*4*) and captured light microscope images of the dorsal abdomen under constant lighting and magnification (3X) on a Nikon SMZ1270 microscope. We then estimated the proportion of white scales on the first abdominal tergite by converting each image to 8-bit grayscale in ImageJ, selecting the region of interest, and calculating the area with brightness values above 128. Area estimates were only weakly sensitive to the precise cutoff since the white and black scales differ markedly in brightness; we therefore chose a value in the middle of the range. We used the R function *lm* to fit a linear model with logit-transformed scaling proportion as the response variable and population of origin as the predictor. We then used the R package *emmeans* to calculate 95% confidence intervals for each population.

### Whole genome resequencing and variant calling

As part of an ongoing 1200 *Aedes aegypti* genomes project, we extracted gDNA from 480 field-collected mosquitoes using the Chemagic DNA tissue protocol and sequenced them to 15x coverage with PE 151bp reads using the Illumina HiSeqX platform. The sequenced mosquitoes included 397 individuals from 24 sub-Saharan African populations collected for this study, 29 additional individuals from sites in Uganda, Kenya, and Burkina Faso that were not included in the main study, 12 individuals of the domestic form collected in 2009 or 2011 in Rabai, Kenya^10^, 20 individuals from Bangkok, Thailand, 18 individuals from Santarem, Brazil, and 4 *Ae. mascarensis* mosquitoes for use as an outgroup.

We initially mapped all sequence data to the L5 reference genome^38^. We identified and removed close relatives from our sample as follows. First, we generated a matrix of relatedness coefficients using the --relatedness2 subprogram from VCFtools^39^ with a set of randomly selected 109,267 biallelic SNPs (MAF>0.05, >1 read in 90% of individuals) preliminarily called with *bcftools*. Second, we hierarchically clustered the coefficients using the R function *hclust* (method=‘average’). Third, we grouped close relatives using the R function *cutree*, with a relatedness cutoff of 0.05 for African samples (corresponding to first cousin or closer relationships) and 0.2 for non-African populations (corresponding to siblings). The more permissive cutoff was used for non-African populations because they are more inbred/bottlenecked, with many individuals showing cousin-like relationships. Finally, we removed all but one randomly-chosen individual from each group of relatives. This left us with 345 sub-Saharan African *Ae. aegypti* genomes from our 24 focal study sites, 14 *Ae. aegypti* from other sites in sub-Saharan Africa, 30 *Ae. aegypti* genomes from outside continental Africa and 4 *Ae. mascarensis* genomes. Most relatives came from the same ovitrap (we sequenced more than one individual from a single ovitrap when ovitrap/egg limited). A smaller number came from nearby ovitraps in the same general location. The 14 *Ae. aegypti* genomes from non-focal sites were used for variant discovery and included in ADMIXTURE and PCA-based analyses (see below) in order to ensure we were sampling as much diversity as possible, but they are not plotted in figures.

We then used three iterative mapping steps to construct an updated African reference based on data from a geographically distributed (Africa only) set of 100 unrelated male mosquitoes (Extended Data Fig. 9). We chose to use males for the update because the L5 reference was constructed using data from males. In each of three iterative mapping steps, we (1) mapped sequence data from the mapping set to the reference using *bwa mem* (MAPQ cutoff of 10), (2) called consensus biallelic SNP genotypes using *bcftools* (*“bcftools mpileup -BI* | *bcftools call -vmOu* | *bcftools view -v snps -q 0.5:alt1* | *bcftools norm -Ou -m -* | *bcftools norm -Oz -d snps”*), and (3) substituted the consensus base into our reference sequence using *bcftools consensus*^40,41^. We used PicardTools^42^ to characterize read mapping quality after each iteration on a set of individuals not used for alternate reference construction (20 individuals; male-female pairs from 10 African populations) (Extended Data Fig. 9). We used a permissive MAPQ cutoff of 10 for the mapping steps because analyses suggested that high levels of sequence divergence from the L5 reference were disrupting initial alignments (Extended Data Fig. 9a-d). Finally, we remapped data from all 480 genomes to the updated third-iteration reference; this included non-African and outgroup samples, which also mapped well to the updated reference (Extended Data Fig. 9e-f). Males and females mapped similarly, except in the region around the sex-determining M-locus (Extended Data Fig. 9g-h). After remapping, we realigned reads near insertions and deletions to improve variant discovery in these regions using GATK IndelRealigner^43^.

We took two different approaches to variant calling - both of which were confined to regions of the genome we inferred to be non-repetitive (repeat masked using the RepeatMasker intervals from the L5 genome) and single copy (mean coverage between 5-30X across individuals). We used the program *ANGSD*^44^ to calculate population-level allele frequencies and genetic diversity, as well as to carry out genotype-likelihood based analyses (see below) for 161,713,099 biallelic single nucleotide polymorphisms (SNPs, *P*<10^−6^). We also called individual genotypes for a filtered set of 14,045,728 high quality biallelic SNPs using bcftools. These SNPs were filtered for coverage across the entire sample (covered by at least 1 read in 90% of individuals) and then called for any individual with sample depth > 8 reads and genotype quality score >30. After individual genotype calling we implemented a further filter for the fraction of individuals genotyped (>75% at any given SNP) and minor allele frequency (MAF>1%). We used the same permissive MAPQ cutoff of 10 for variant calling as used for generating the updated reference genome (see above) in order to minimize potential problems with aligning alternate haplotypes. Note that our additional MAF and genotyping filters help protect against SNP calls from false positive alignments. Hard genotype calls (or subsets thereof) were used for *ADMIXTURE*, principal components analyses (PCA), *PCAdapt*, and *Dsuite* analyses (see below).

### Population structure analyses

We characterized population structure using two alternative approaches based on a set of 1,000,000 unlinked SNPs selected in *PLINK* (step size 100, cutoff 0.1, --thin-count 1000000)^45^. First, we used *ADMIXTURE*^21^ to assign individuals to variable numbers of population clusters for K=2-10, with K=3 minimizing cross-validation error (Extended Data Fig. 3b). Second, we used *PLINK* to carry out principal components analysis (PCA). One sample from Ngoye, Senegal (NGO) was a clear outlier in ancestry, showing strong affinity with West African generalist populations while all other individuals from this population showed consistent affiliation with human specialists (Fig. 3c-d); we excluded this putative recent migrant from subsequent *F*_*ST*_, *PBS, ABBA-BABA, π*, and *d*_*xy*_ analyses involving Ngoye (see below).

### Gene flow and divergence analyses

We used *ANGSD* (subprogram *realSFS*) to calculate pairwise *F*_*ST*_ between populations and a custom script to turn these FST values into the Population Branch Statistic (*PBS*, essentially polarized *F*_*ST*_) for NGO, THI, OGD, KED, and OHI, using BTT as a nearby generalist reference population and FCV as an outgroup^22^. Using alternative reference and outgroup populations yielded similar results. We used a permutation testing approach to construct a distribution of 5Mb *PBS* values under the null hypothesis of homogeneous differentiation across the genome. More specifically we shuffled the genomic locations of 10kb windows (which preserves local linkage patterns) with the constraint that shuffling could only occur among regions matched for average genetic diversity (average of BTT and focal population). We then used this null distribution to identify ≥5Mb regions of elevated *PBS* at a false discovery rate of 0.05 using a two-tailed test. The choice of a 5Mb window restricts us to largish regions that are either relatively new (giving recombination limited time to break up divergent chromosomal regions) or contain several tightly linked loci. However, using smaller regions (e.g. 1Mb or 100kb) identified similar patterns.

We used *ABBA-BABA*-related statistics to further explore patterns of divergence between populations. These statistics test for an excess of shared derived variation between lineages in order to distinguish gene flow from the incomplete lineage sorting (ILS) that can occur during a simple tree-like branching process. For more on expected genome-wide and locus-specific patterns of derived allele sharing under ILS and gene flow, see^23^. First, we used *Dsuite*^46^ to confirm that the populations in our dataset did not conform to the strict tree-like model, which is expected since all populations belong to the same species and almost certainly exchange genes. Indeed, we strongly rejected the null tree-like hypothesis (block-jacknife *P*<10^−7^) for all three-population trees with *Ae. mascarensis* as an outgroup.

We then explored potential heterogeneity in gene flow across the genome using the *f*_*D*_ statistic (calculated in 5Mb windows with a 10kb step). The *f*_*D*_ statistic uses shared derived genetic variation to estimate the fraction of ancestry at a specific locus derived from gene flow between branches in a specified tree^23^. We calculated *f*_*D*_ from *ANGSD* population allele frequencies using a custom python script for the tree (BTT, X; BKK, *mascarensis*) to identify regions of the genome showing elevated levels of shared derived variation between the focal population (X=NGO, THI, or OGD) and non-African human specialists (BKK). We do not think such shared derived variation is necessarily derived from introgression back to Africa from non-African populations. Instead, it may reflect relationships present in ancestral populations, before *Ae. aegypti* left Africa. Differentiating between these two hypotheses is out-of-scope for this study but will be addressed in a future study incorporating much more genomic data from outside Africa. Regardless, we expect shared derived variation between human-preferring populations within and outside Africa to be present in regions that code for human-adaptive traits and thus experience reduced gene flow between human- and animal-preferring populations within Africa. We used Fisher’s exact test to test whether the top 10% of non-overlapping 5Mb *f*_*D*_ windows were significantly enriched in our *PBS* outlier regions for each focal population.

To help interpret measures of between-population divergence (i.e. *F*_*ST*_ and *PBS*), we used *ANGSD* to estimate levels of genetic diversity (*π*) across the genome for each population and the perl script getDxy.pl (modified to skip variant sites not covered in one population) from ngsTools^47^ to calculate *d*_*xy*_ (Extended Data Fig. 5). We also calculated normalized *d*_*xy*_ (Extended Data Fig. 5c) by dividing *d*_*xy*_ for a given population pair by mean *d*_*xy*_ for all pairs of NGO, THI, OGD, BTT, and FCV. We calculated normalized *π* (Extended Data Fig. 5c) by dividing population *π* by mean *π* across all populations.

We used PCAdapt^25^ to test whether specific SNPs were associated with specialist ancestry across the subset of high-quality, biallelic SNPs from our hard-called set that had minor allele frequency >0.05 (n=5,369,564 SNPs) (PCAdapt parameters: K=3, method “componentwise”, and LD clumping with a size of 200 and a cutoff of 0.1).

### Climate and population projections

We predicted future changes in host odor preference at each sampling location by plugging climate and human population density change projections for 2050 into our final, fitted, ecological model – including human population density (calculated within 20km radius), precipitation seasonality (Bio15, third degree monotonic polynomial) and precipitation in the warmest quarter (Bio18, linear).

Climate change projection data came from the Coupled Model Intercomparison Project Phase 5 (CMIP5) based on Representative Concentration Pathway 8.5 (RCP8.5) scenarios^27^. RCP8.5 is considered the business-as-usual scenario for future greenhouse gas concentrations, reflecting minimal mitigation efforts. The CMIP5 effort contributed to the International Panel on Climate Change (IPCC) Fifth Assessment Report. A new modeling effort, CMIP6, is currently underway but complete data are not yet available. Projected climate data from a global climate model (GCM) cannot be directly compared to present-day observational climate data due to model biases and measurement error. Failing to account for these biases can result in misinterpreting structural differences between the two datasets as potential climate change effects. Instead, the projection data must first be bias-corrected by calculating the relative or absolute change between current and future climates for the variable of interest, using solely the GCM output. This relative or absolute change can then be applied to observational data. Depending on the resolution of the observational climate data, projections may also be downscaled – i.e. the resolution of the model output improved. The Worldclim projection data has undergone both downscaling and bias-correction processes such that it can be compared with the observational data used in our present-day analysis. Absolute changes were used for temperature and relative changes were used for precipitation. Further details of these processes are available at https://www.worldclim.org/downscaling. There is relatively high agreement across models in terms of the spatial distribution of projected Bio15 and Bio18 changes (Extended Data Fig. 7).

Population projection data came from the United Nations medium-variant scenario^48^. This scenario assumes existing high fertility populations will experience a fertility decline over the coming century, as economic development increases. Despite falling fertility, sub-Saharan Africa is expected to see an increase in the total number of births over the next several decades relative to the recent past. High birth numbers coupled with increasing life expectancy will lead to 1.05 billion increase in population in sub-Saharan African countries by 2050, 52% of the additional global population in this timeline^49^. We used urban and rural, medium-variant projections for each country and calculated growth rates by comparing 2050 numbers with those from 2015. Note, our field collections were conducted between 2015 and 2018 (mostly 2017-2018) making 2015 numbers more applicable than any other available estimates. We then applied these growth rates to the baseline population data for each location. Urban and rural locations were considered separately because urban populations are expected to grow at a faster rate than rural populations over this time period. Urban populations were defined as those with current population density > 400 humans/km^2^ calculated with a 20km buffer (Extended Data Table 1). These populations were all from areas that we observed to be dominated by human structures and activities (Extended Data Table 1). A few intermediate density locations fell below this cutoff and were classified as rural. More specifically, KWA, OHI and NGO are rural towns, while ABK and SHM are wild areas on the far outskirts of what most would consider urban areas (densities 173-288 humans/km^2^; Extended Data Table 1). The other sites classified as rural had much lower densities (1-67 humans/km^2^; Extended Data Table 1).

## Data and code availability

Raw genomic data are deposited in the NCBI SRA under the accession code PRJNA602495 and will be made available upon publication. Other raw data and scripts are available at github.com/noahrose.

## Acknowledgements

The authors thank Siyang Xia, Christophe Paupy, and Bryan Grenfell for valuable feedback on early results, Boy Ponlawat for generously providing mosquito eggs used to establish a laboratory reference colony from Thailand, Francis Mulwa, Gilbert Rotich, Gilbert Bianquinche, Marc F. Ngangué for field assistance, the National Park Services and rangers of Kenya, Gabon, and Ghana for providing access to forest areas, and a large number of local residents who gave advice and assistance at all field sites. The findings and conclusions in this report are those of the authors and do not necessarily represent the official position of CDC.

## Author contributions

N.H.R., M.S., A.B., J.L., D.A., O.B.A., N.I., S.O., J.A., J.P.M., R.S., and C.S.M. planned and carried out field collections. N.H.R., A.L.K., M.S., and J.R.P. established laboratory colonies. N.H.R, N.I., and A.L.K carried out behavioral experiments. N.H.R. and E.G.E. carried out morphological quantification. B.J.W. and J.E.C. carried out genome sequencing and alignment. J.R.P. and A.G.S. provided non-African samples for sequencing. N.H.R., B.J.W., J.E.C. and C.S.M analyzed genomic data. R.E.B. modeled future conditions. N.H.R. and C.S.M. conceived the project and wrote the manuscript, with input at all stages from many other authors.

## Extended Data

**Extended Data Fig. 1.**
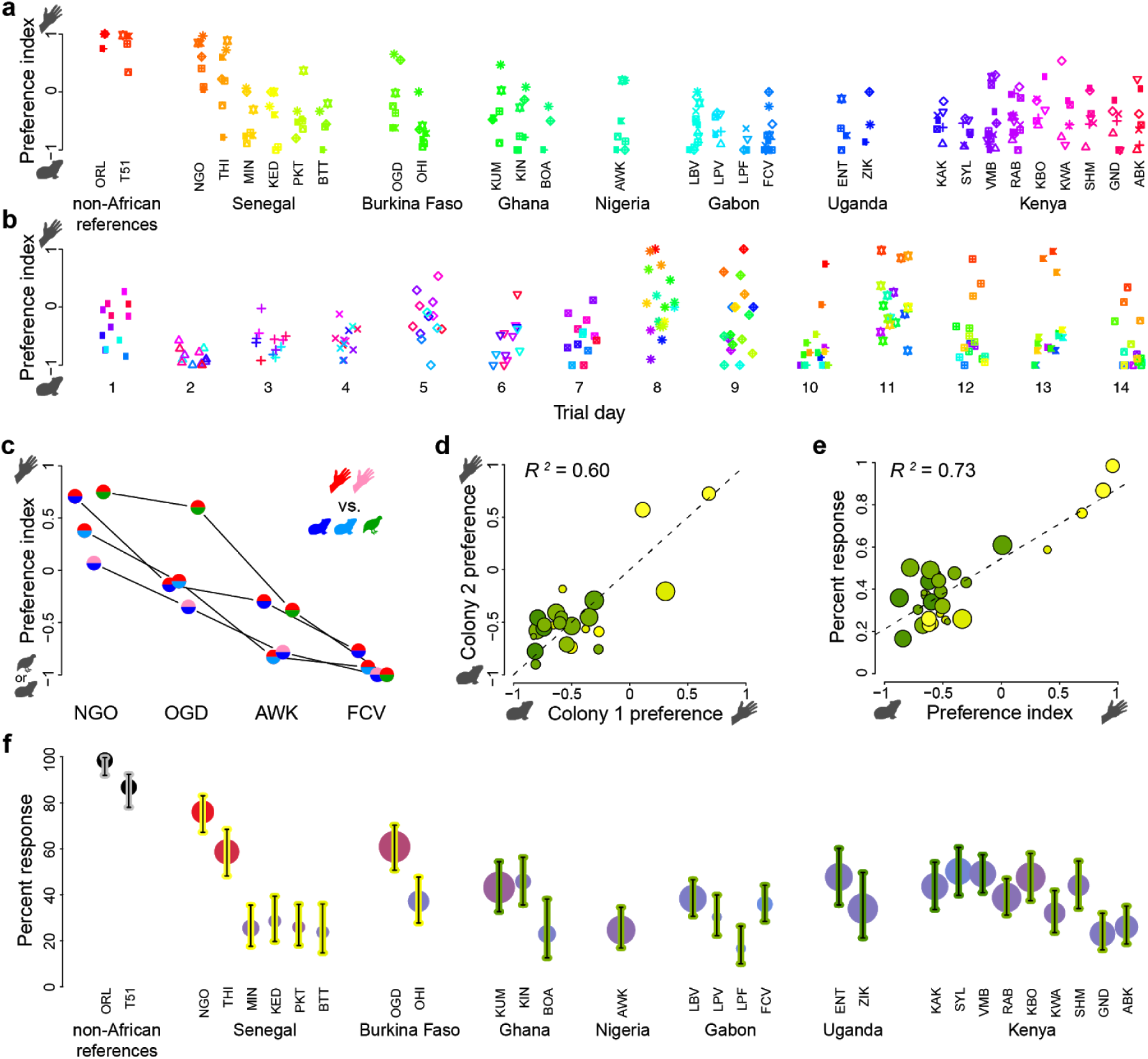
Additional details for host preference assays. **a**-**b**, Raw preference indices for all trials plotted by location (a) or trial day (b). Color indicates location and symbols indicate trial day. **c**, Preference estimates for four populations derived from tests with alternative human subjects (top color of each circle) and animal species (bottom color of each circle) (n=3-8 trials, each including 37-110 females per colony/host combination). Data for one human versus either of two guinea pigs (red/light blue and red/dark blue) come from same experiments summarized in Fig. 1d. Data for same human versus quail (red/green) and second human versus guinea pig (pink/dark blue) are new. **d**, Host preference was strongly correlated for replicate colonies from the same location (Pearson *R*^*2*^=0.60, *P*=1.5×10^−5^). **e**, Preference and overall response rate (rate of choosing either host) were strongly correlated across locations (*R*^*2*^=0.73, *P*=4.4×10^−9^). **f**, Overall response rates for each location plotted and analyzed as in Fig. 1d.

**Extended Data Fig. 2.**
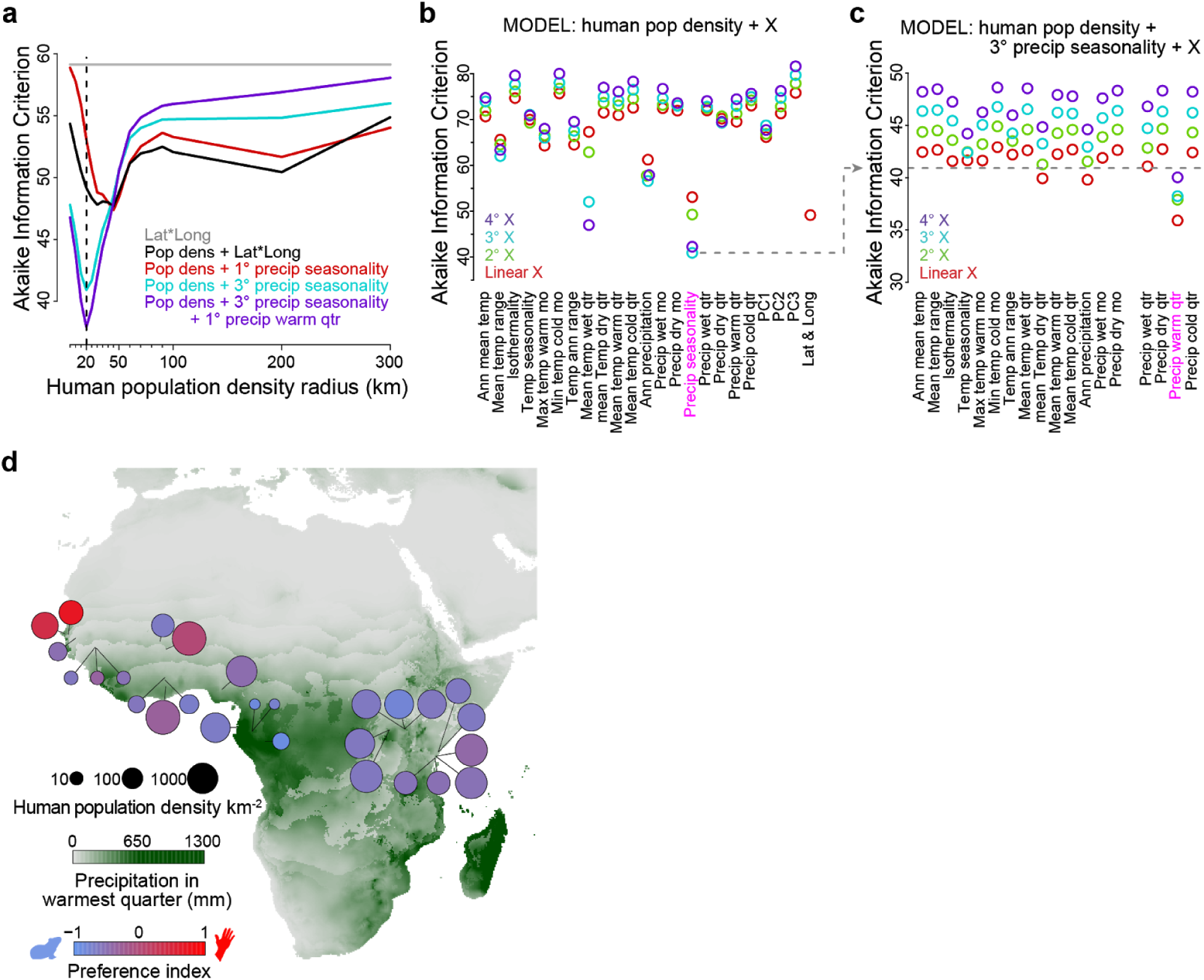
Additional details for modelling effects of human population density and climate on host preference. **a**, Akaike Information Criterion (AIC) for linear models of host preference as a function of various combinations of region (latitude/longitude), specific climate variables, and human population density estimated across different scales (x-axis). All models incorporating population density perform better than a null model that includes region only (Lat*Long, grey line). Dotted vertical line highlights the spatial scale at which population density best predicted behavior in our final model (purple). **b**, AIC for linear models of host preference as a function of human population density plus individual bioclimate variables. Precipitation seasonality (pink) provided the best fit, especially when modeled as a monotonic cubic polynomial. **c**, After removing the effects of population density and precipitation seasonality, a linear effect of precipitation in the warmest quarter (pink) further decreases AIC. No further variable improves the model after including this second climate effect. **d**, Map of collection localities equivalent to Fig. 1c except on background map of variation in precipitation in the warmest quarter (rather than precipitation seasonality).

**Extended Data Fig. 3.**
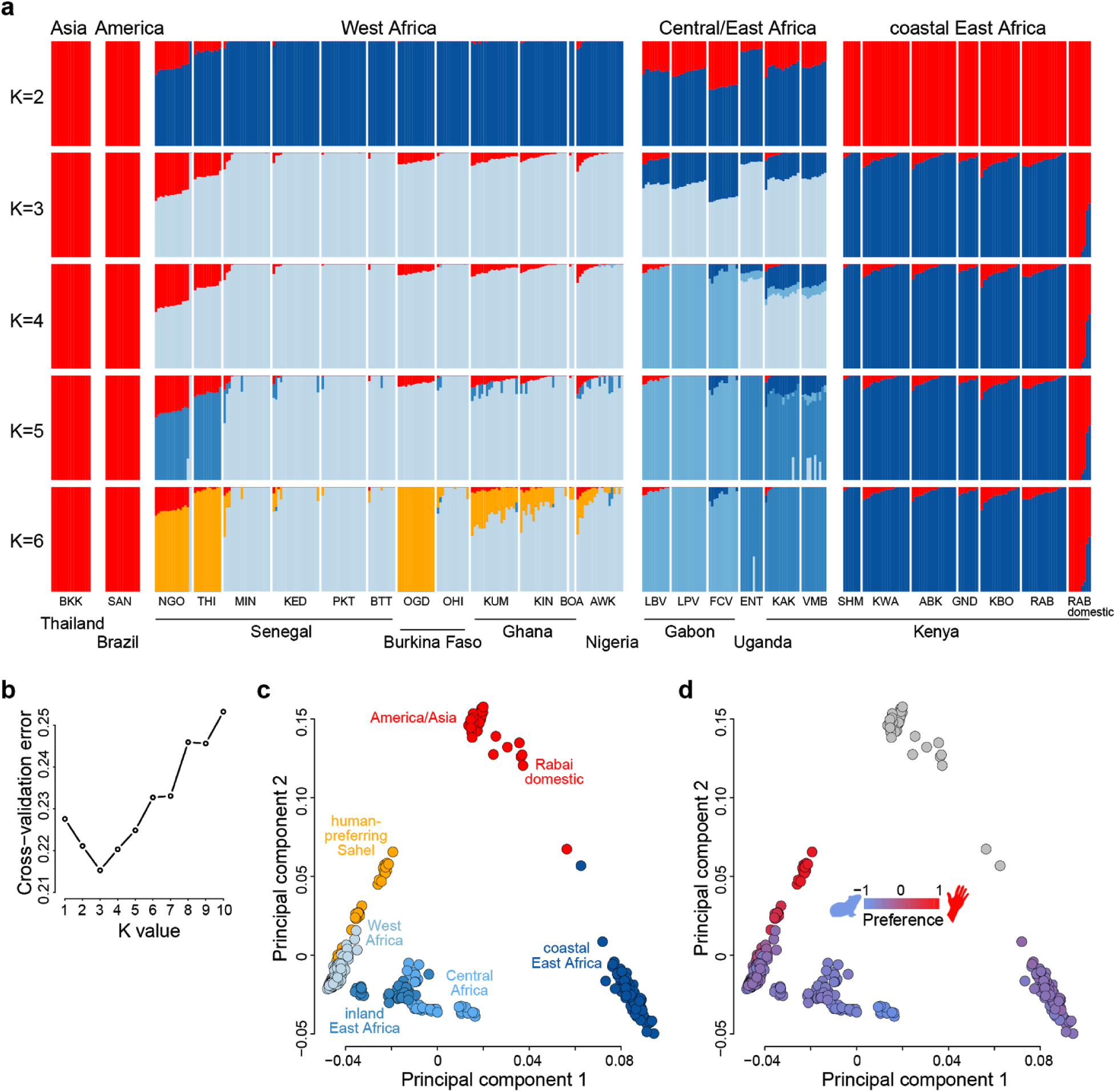
Additional details for analyses of population structure. **a**, Ancestry proportions from ADMIXTURE analysis for K=2-6. **b**, Cross-validation error for analysis in (a) is minimized for K=3. **c**-**d**, Principal components analysis shows three main clusters, with sub-structure corresponding to the clusters found at higher K values (c) and, remarkably, to behavior (d). The non-African and Rabai domestic samples are shown in grey in panel (d) because their behavior was not tested in this study. However, we expect them to be strongly human-preferring based on previous work ^4^ as well as testing of the T51 colony (from Thailand) and ORL colony (likely of largely American descent) in this study (Fig. 1d).

**Extended Data Fig. 4.**
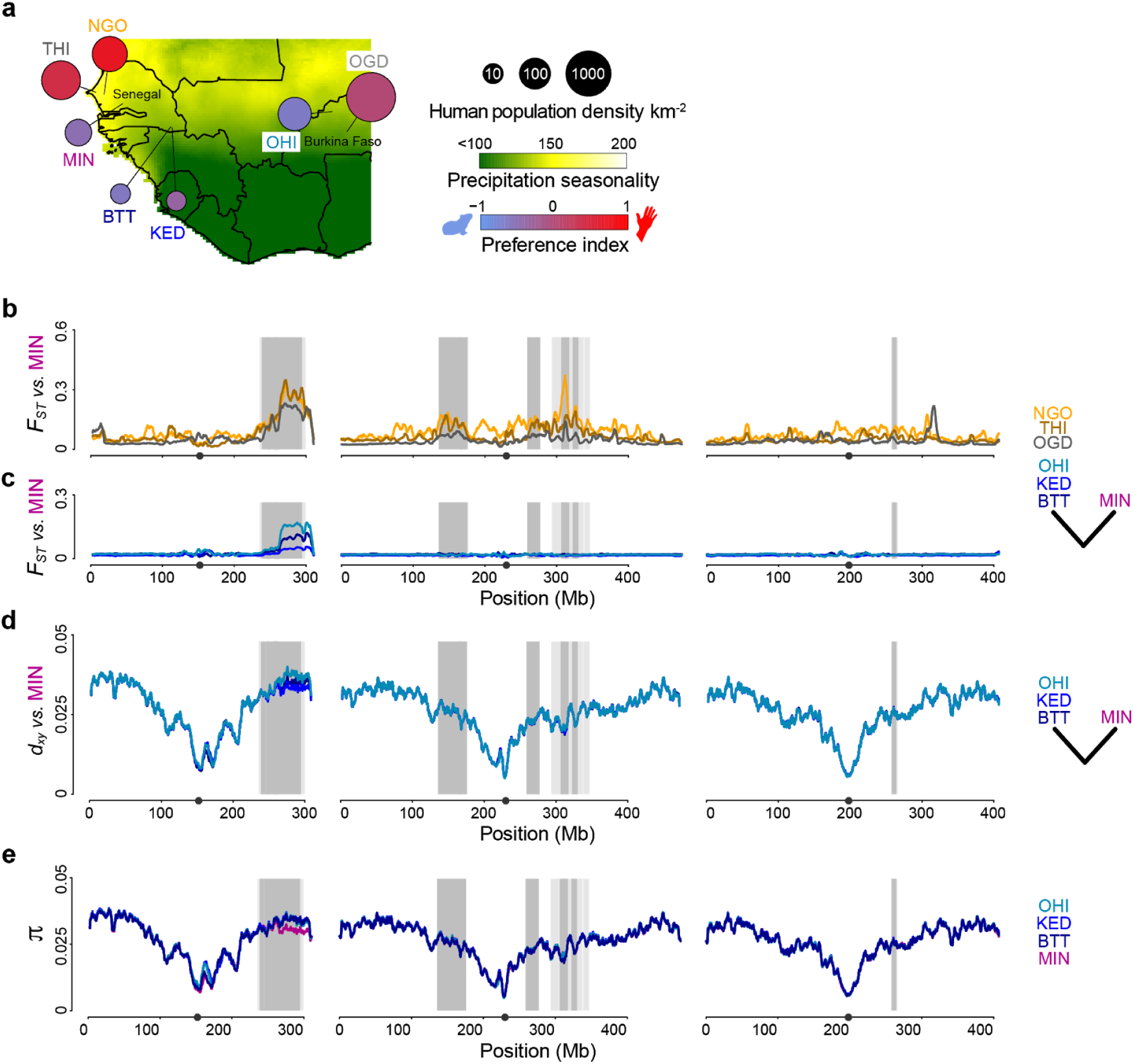
Mindin, Senegal (MIN) shows signs of a selective sweep and/or structural rearrangement in the chromosome 1 outlier region. **a**, Map of sites addressed in this figure (taken from Fig. 1c with precipitation seasonality scale adjusted to better illustrate climate gradient in Senegal and Burkina Faso. Mindin is an animal-seeking population at the bleeding edge of the geographic transition to human-seeking in the Sahel region of Senegal. **b**, Mosquitoes from Mindin show elevated divergence from human-seeking populations at the distal end of chromosome 1, as do other animal-seeking populations (e.g. BTT; Extended Data Fig. 5b). Grey shading indicates *PBS* outlier regions. **c**, Unexpectedly, mosquitoes from Mindin also show step-like, elevated divergence in this chromosomal region from other animal-seeking populations, despite having similar host preference. Elevated divergence in (c) is a function of variably elevated absolute differentiation (*d*_*xy*_) (**d**) and markedly reduced genetic diversity (*π*) in Mindin (**e**).

**Extended Data Fig. 5.**
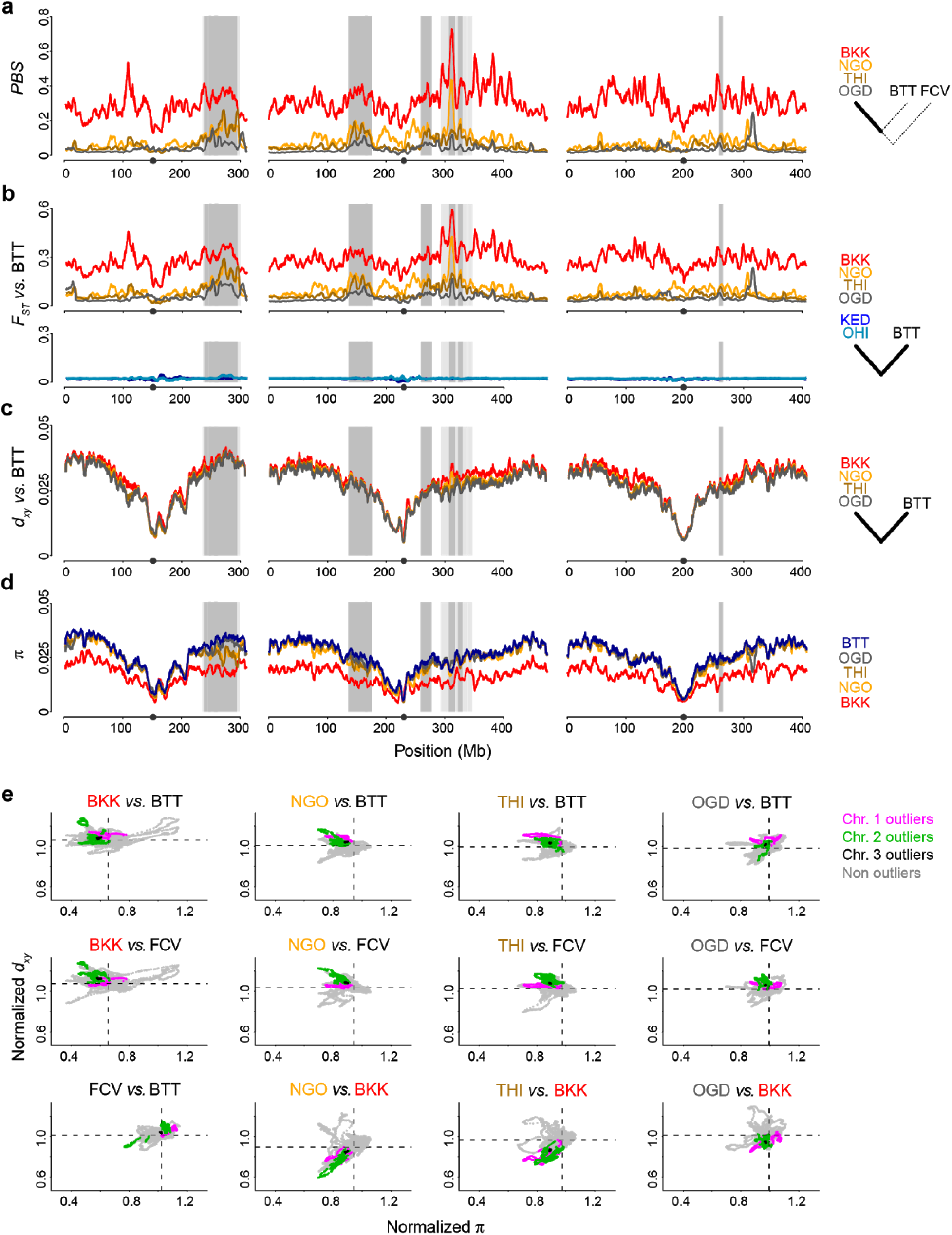
Additional details for analyses of genomic divergence between human- and animal-seeking populations. **a**, Polarized genomic divergence (Population Branch Statistic, *PBS*) as shown in Fig. 3f with the addition of a human specialist population from outside Africa (Bangkok, BKK). BKK shows elevated differentiation across the genome. **b**, Non-polarized divergence (*F*_*ST*_) from the animal-seeking population Bantata (BTT). Human-seeking populations show elevated differentiation in *PBS* outlier regions. Animal-seeking populations show minimal differentiation even when geographically distant from BTT (e.g. Ouahigouya, OHI). BKK again shows higher differentiation across the genome. Patterns in (b) are driven by variation in (**c**) absolute genetic differentiation (*d*_*xy*_) and (**d**) within-population diversity (*π*) in the same populations. **e**, Analyses of normalized *d*_*xy*_ and normalized *π* indicate that human-seeking populations have both elevated absolute divergence and reduced diversity within *PBS* outlier regions (pink, green, black), suggesting that both sweeps and subsequent selection against gene flow have driven divergence in these regions. Conversely, human-seeking populations inside Africa show reduced levels of absolute divergence from the non-African population (BKK) in these regions (last three subpanels), consistent with shorter coalescence times and a common origin.

**Extended Data Fig. 6.**
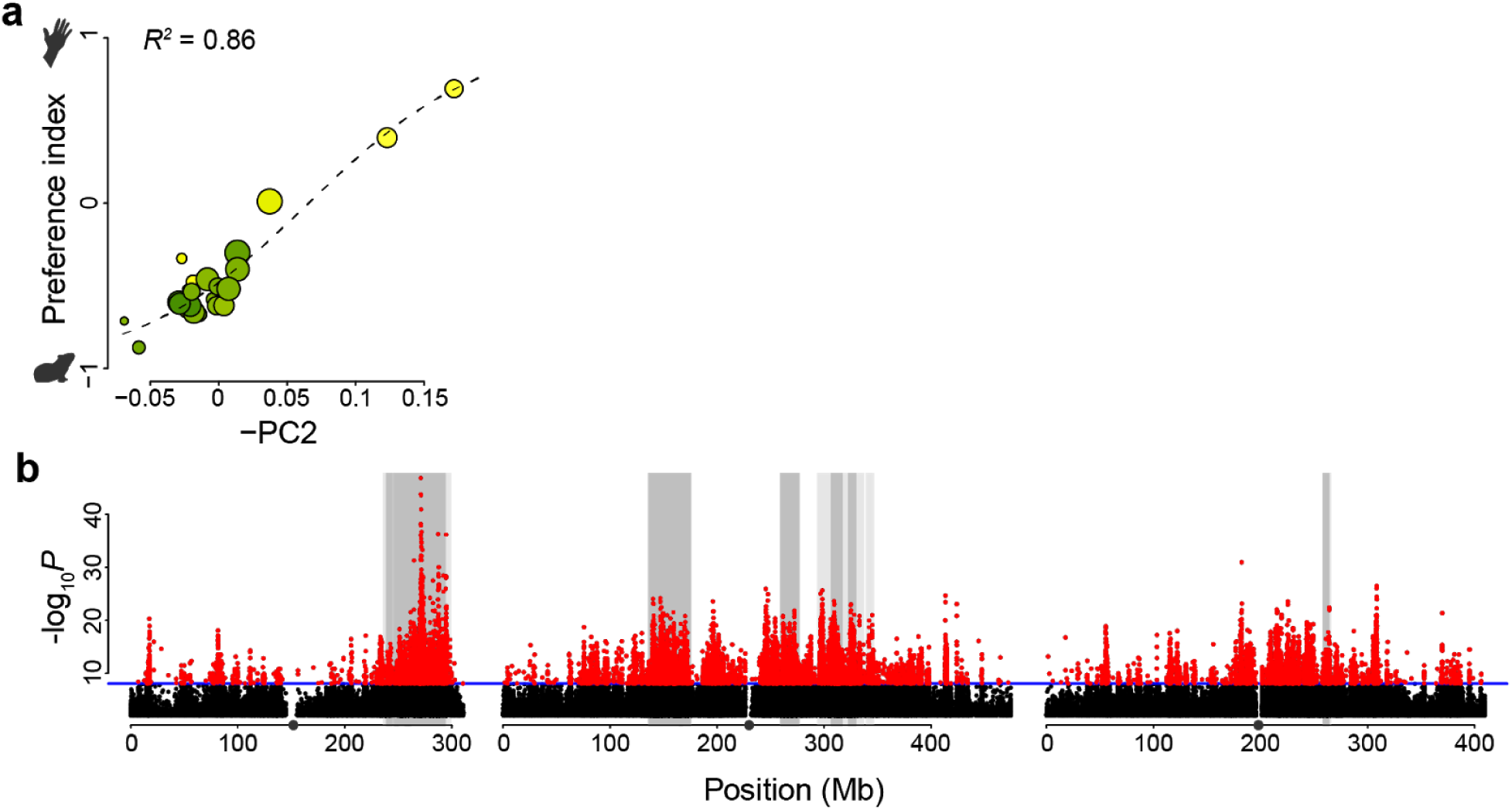
PCAdapt analysis identifies genome-wide variants associated with human specialist ecology. **a**, The second principle component from a PCAdapt analysis of African genomes shows a striking correlation with host preference. This result is similar to the outcome of the principal components analysis of population structure shown in Extended Data Fig. 3d, where the second PC is clearly also strongly correlated with behavior within Africa. **b**, Thousands of SNPs were more strongly associated with PC2 than expected under neutral evolution, suggesting positive selection (n=16,782 SNPs in red, Bonferroni-adjusted *P*<0.05). Consistent with analyses of genomic divergence between human- and animal-preferring populations in West Africa (Fig. 3f), the most significant SNPs are concentrated in *PBS* outlier regions (grey shading). Points with unadjusted *P*>0.01 not plotted.

**Extended Data Fig. 7.**
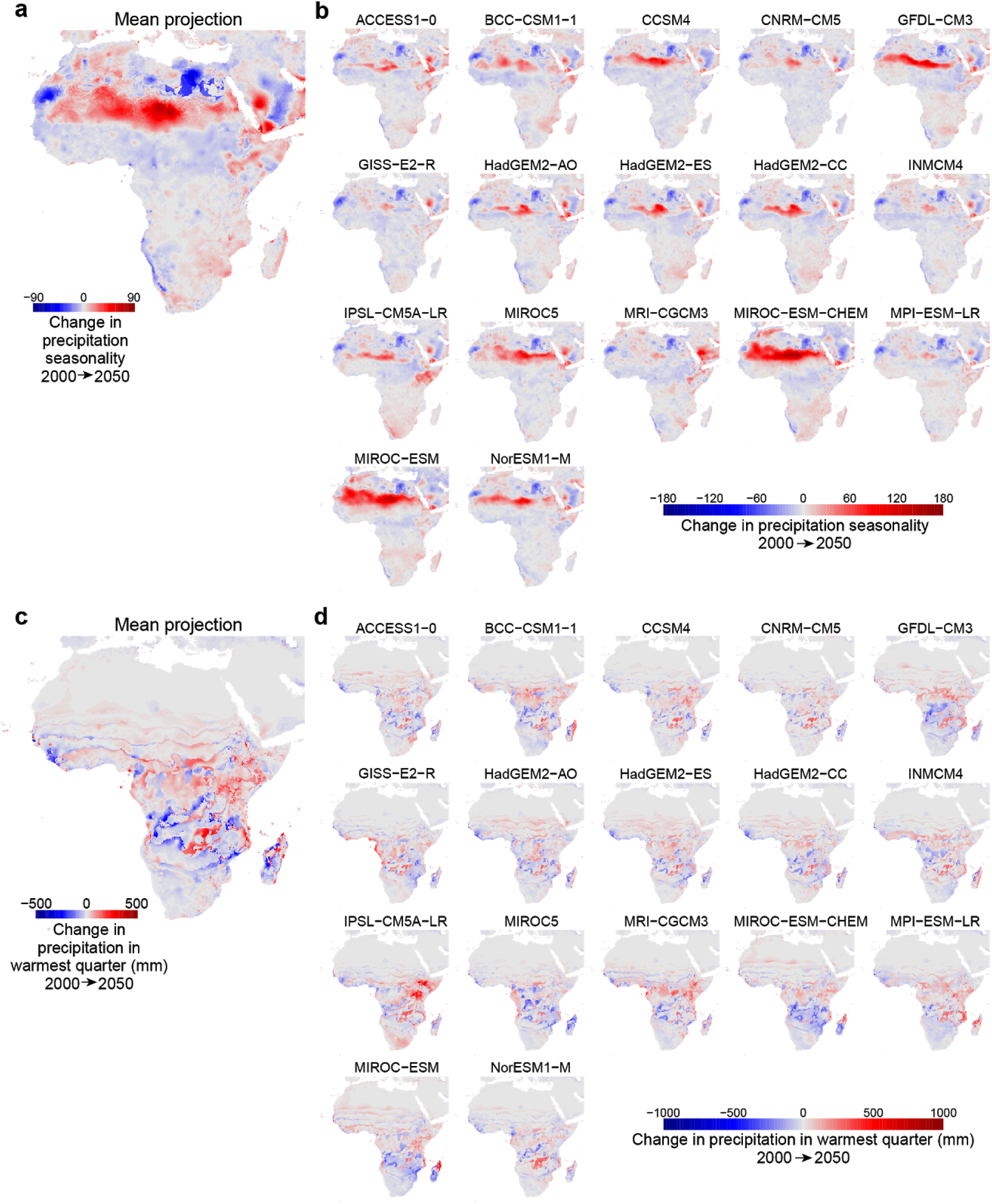
Projected change in preference-related climate variables from 2000 to 2050. **a**, Mean projected change in precipitation seasonality across all climate models. **b**, Individual model projections for precipitation seasonality. Note that scale differs from panel A. **c**-**d**, same as a-b except for precipitation in the warmest quarter.

**Extended Data Fig. 8.**
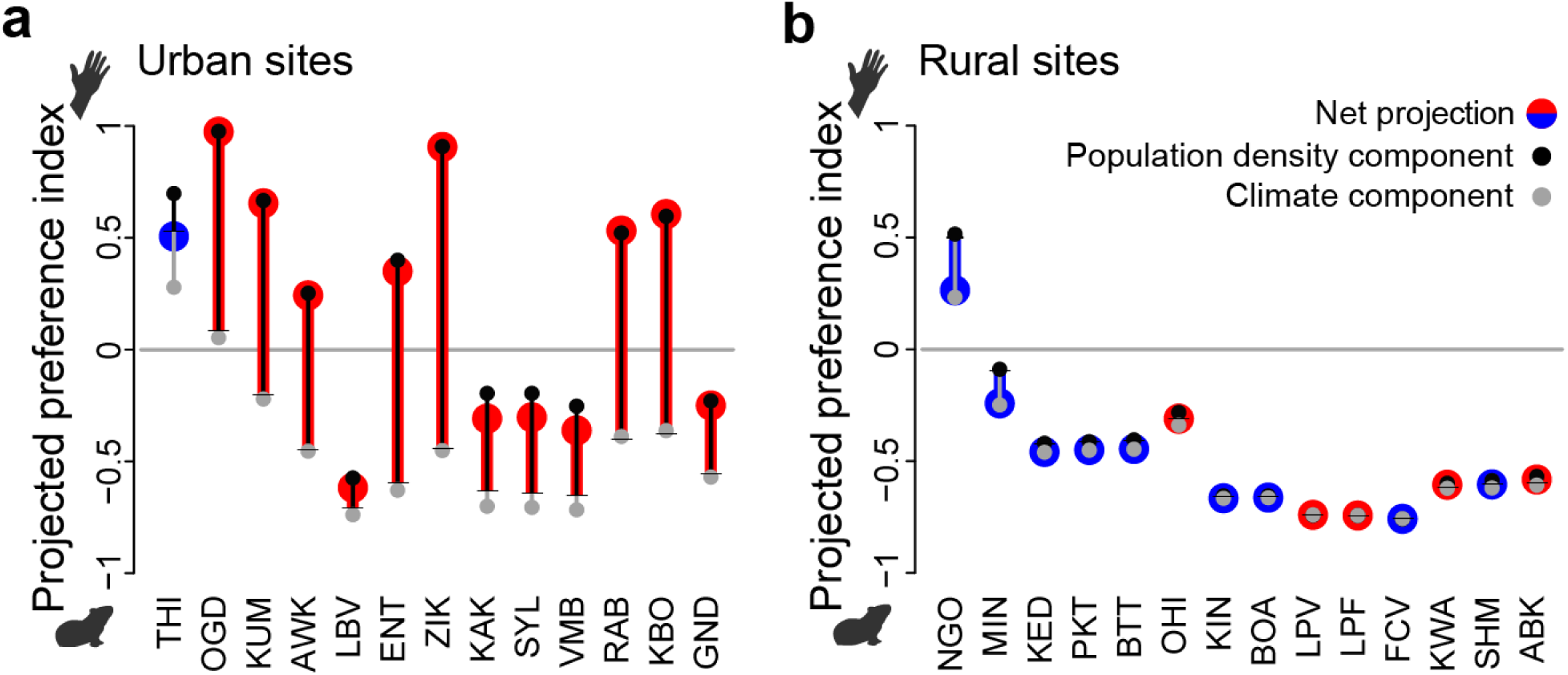
Additional details on host preference projections for 2050. Expected effects of changes in human population density (black) and climate (grey) are shown separately. Net effects are shown in red (increases) or blue (decreases). Note that publicly available human population growth rate projections are different for urban (**a**) and rural (**b**) sites. We classified sites as urban if current population density exceeded 300 humans km^-2^.

**Extended Data Fig. 9.**
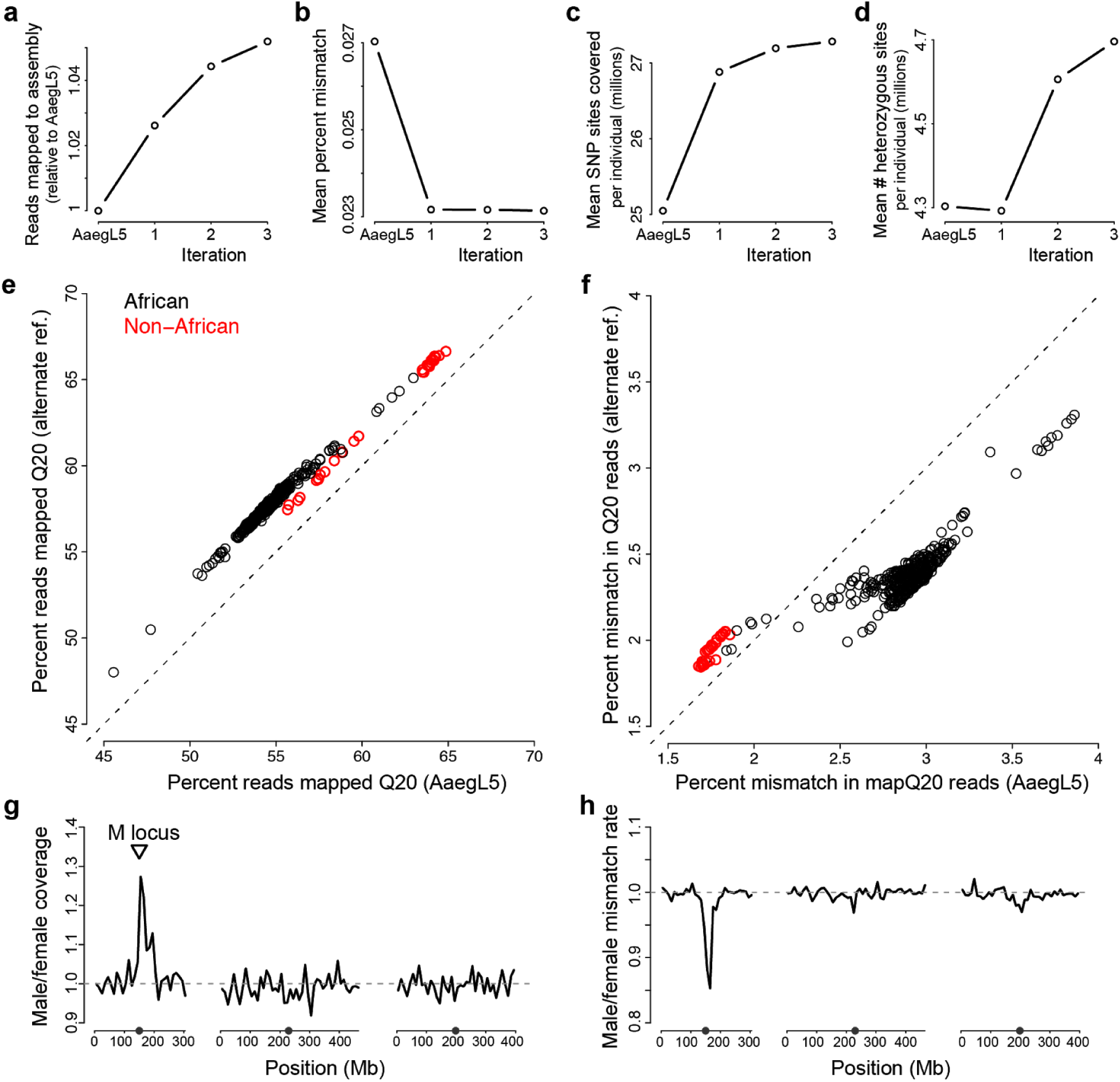
Construction of an alternate reference genome improved read mapping and variant discovery. **a**-**d**, Comparison of four mapping statistics across three iterative updates to the AaegL5 genome for a panel of 20 individuals (one male and one female from each of 10 African populations): **a**, Total number of reads mapped with MAPQ>20. **b**, fraction of mismatched bases. **c**, mean number of polymorphic sites per individual covered by at least 10 reads with MAPQ>10 (polymorphic sites defined as those with at least 5 alternate allele reads present among summed reads from all samples). **d**, average number of heterozygous sites per individual covered by >10 reads with MAPQ>10. **e**-**f**, Improvements in high quality mapping rates (MAPQ>20) (e) and mismatch rates (f) were most apparent for African mosquitoes (black). Use of the alternate reference also improved high quality mapping rates for non-African mosquitoes (red, e), but increased mismatch rates (f). **g**-**h**, Males and females had similar coverage (g) and mismatch rates (h) when mapped to the alternate reference, except around the sex-determining M locus. At the M locus, males had higher coverage and fewer mismatches. Note the AaegL5 assembly and alternate reference were both constructed using male mosquitoes. In panel h, the pair from Franceville, Gabon (FCV) is excluded due to a male outlier showing elevated genome-wide mismatch rates.

**Extended Data Table 1.** Details of collection localities. Climate variables and population densities are calculated within a 20km radius. Latitude and Longitude are the average coordinates of egg-positive ovitraps from collections.

| Location | Country | Code | Latitude | Longitude | Habitat | Human Population Density km<sup>-2</sup> | Precip. Seasonality (coef. var.) | Precip., Warmest Qtr. (mm) | Pref. Index |
| --- | --- | --- | --- | --- | --- | --- | --- | --- | --- |
| Ngoye | Senegal | NGO | 14.63960875 | -16.43174115 | Town | 232.67 | 157.76 | 135.71 | 0.69 |
| Thies | Senegal | THI | 14.75784115 | -16.93863845 | Town | 421.12 | 159.80 | 325.08 | 0.40 |
| Mindin | Senegal | MIN | 14.07473585 | -15.28339615 | Town | 53.61 | 140.43 | 59.95 | -0.48 |
| Kedougou | Senegal | KED | 12.55867615 | -12.18635415 | Town | 12.25 | 124.44 | 54.75 | -0.52 |
| PK10 | Senegal | PKT | 12.60970505 | -12.2474927 | Forest | 12.29 | 124.94 | 52.50 | -0.33 |
| Bantata | Senegal | BTT | 12.6749906 | -12.3270609 | Forest | 12.29 | 125.46 | 49.88 | -0.59 |
| Ouaga-dougou | Burkina Faso | OGD | 12.37455556 | -1.49805556 | Town | 2153.80 | 119.76 | 108.72 | 0.01 |
| Ouahi-gouya | Burkina Faso | OHI | 13.583331 | -2.416665 | Town | 172.60 | 130.56 | 114.35 | -0.64 |
| Kumasi | Ghana | KUM | 6.67544745 | -1.5665543 | Town | 2264.55 | 53.99 | 332.28 | -0.30 |
| Kintampo | Ghana | KIN | 8.052798 | -1.725279 | Town | 37.23 | 65.89 | 224.41 | -0.58 |
| Boabeng | Ghana | BOA | 7.708767 | -1.6973721 | Forest | 67.04 | 64.17 | 242.57 | -0.67 |
| Awka | Nigeria | AWK | 6.2509878 | 7.00215995 | Town | 1086.14 | 72.51 | 232.17 | -0.46 |
| Libreville | Gabon | LBV | 0.4259774 | 9.4483769 | Town | 799.57 | 68.76 | 880.57 | -0.66 |
| Lope Village | Gabon | LPV | -0.1050309 | 11.60796015 | Town | 1.27 | 69.40 | 553.49 | -0.71 |
| Lope Forest | Gabon | LPF | -0.204279 | 11.600281 | Forest | 1.27 | 69.95 | 562.14 | -0.84 |
| Franceville | Gabon | FCV | -1.63997525 | 13.582982 | Town | 38.54 | 59.72 | 633.48 | -0.87 |
| Entebbe | Uganda | ENT | 0.06444 | 32.44694 | Town | 881.75 | 41.60 | 357.38 | -0.60 |
| Zika | Uganda | ZIK | 0.122214 | 32.526191 | Town | 1588.71 | 40.20 | 336.43 | -0.60 |
| Kakamega | Kenya | KAK | 0.2845454 | 34.7543336 | Town | 707.17 | 38.87 | 342.16 | -0.62 |
| Shinyalu | Kenya | SYL | 0.22688755 | 34.8133263 | Town | 727.46 | 38.05 | 372.10 | -0.78 |
| Virembe | Kenya | VMB | 0.23893465 | 34.84780885 | Town | 671.64 | 38.34 | 378.52 | -0.61 |
| Rabai | Kenya | RAB | -3.9342022 | 39.5730709 | Town | 1291.36 | 65.53 | 238.85 | -0.52 |
| Kaya Bomu | Kenya | KBO | -3.9308373 | 39.59651005 | Forest | 1392.59 | 66.18 | 239.75 | -0.40 |
| Kwale | Kenya | KWA | -4.1721596 | 39.44844215 | Town | 184.84 | 62.89 | 166.48 | -0.50 |
| Shimba Hills | Kenya | SHM | -4.25009085 | 39.4006515 | Forest | 187.13 | 64.95 | 133.88 | -0.54 |
| Ganda | Kenya | GND | -3.2204869 | 40.06458775 | Town | 504.84 | 80.23 | 181.49 | -0.62 |
| Arabuko | Kenya | ABK | -3.29709435 | 39.94876745 | Forest | 288.57 | 71.83 | 172.13 | -0.62 |

**Extended Data Table 2.**
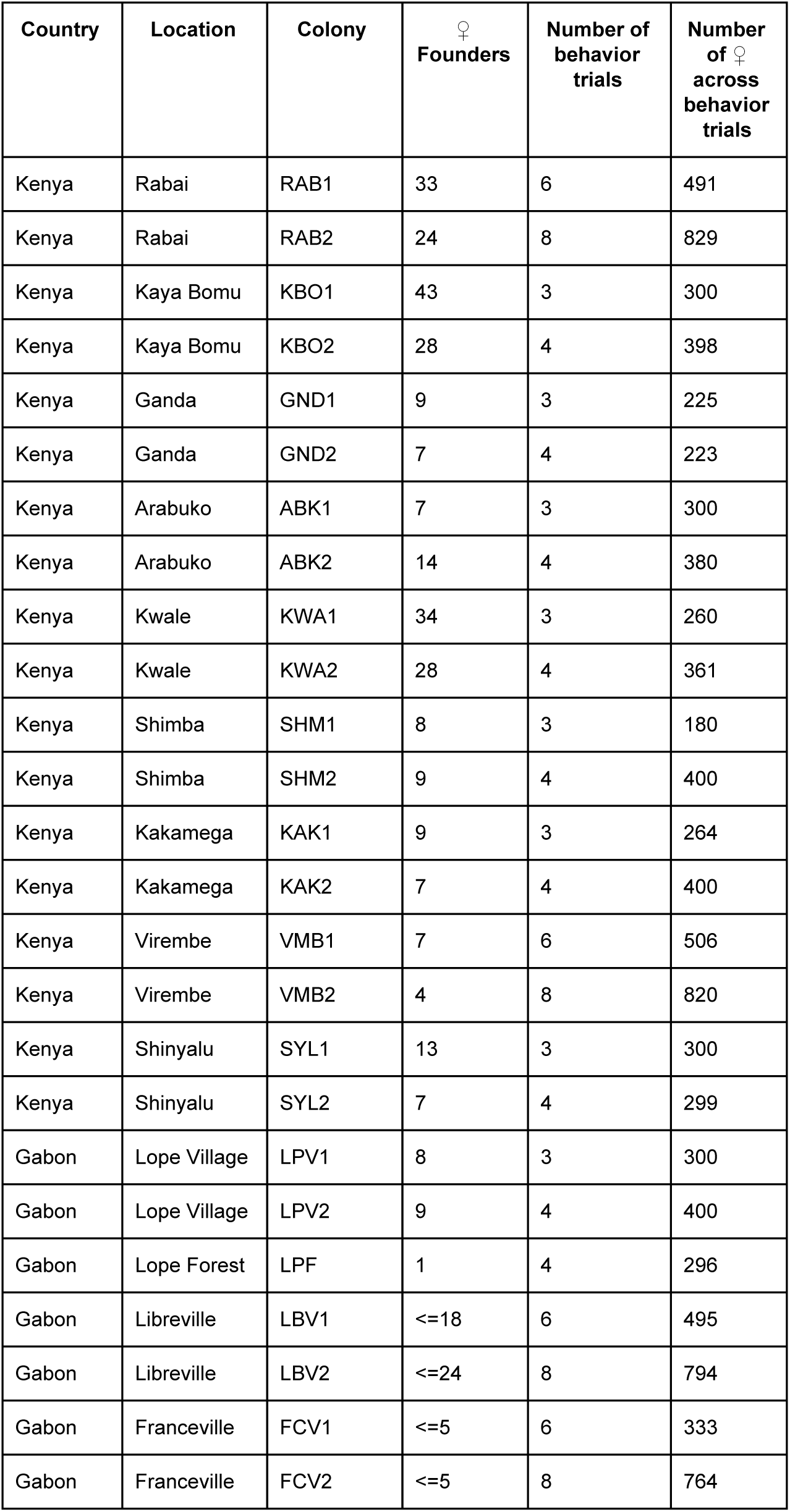
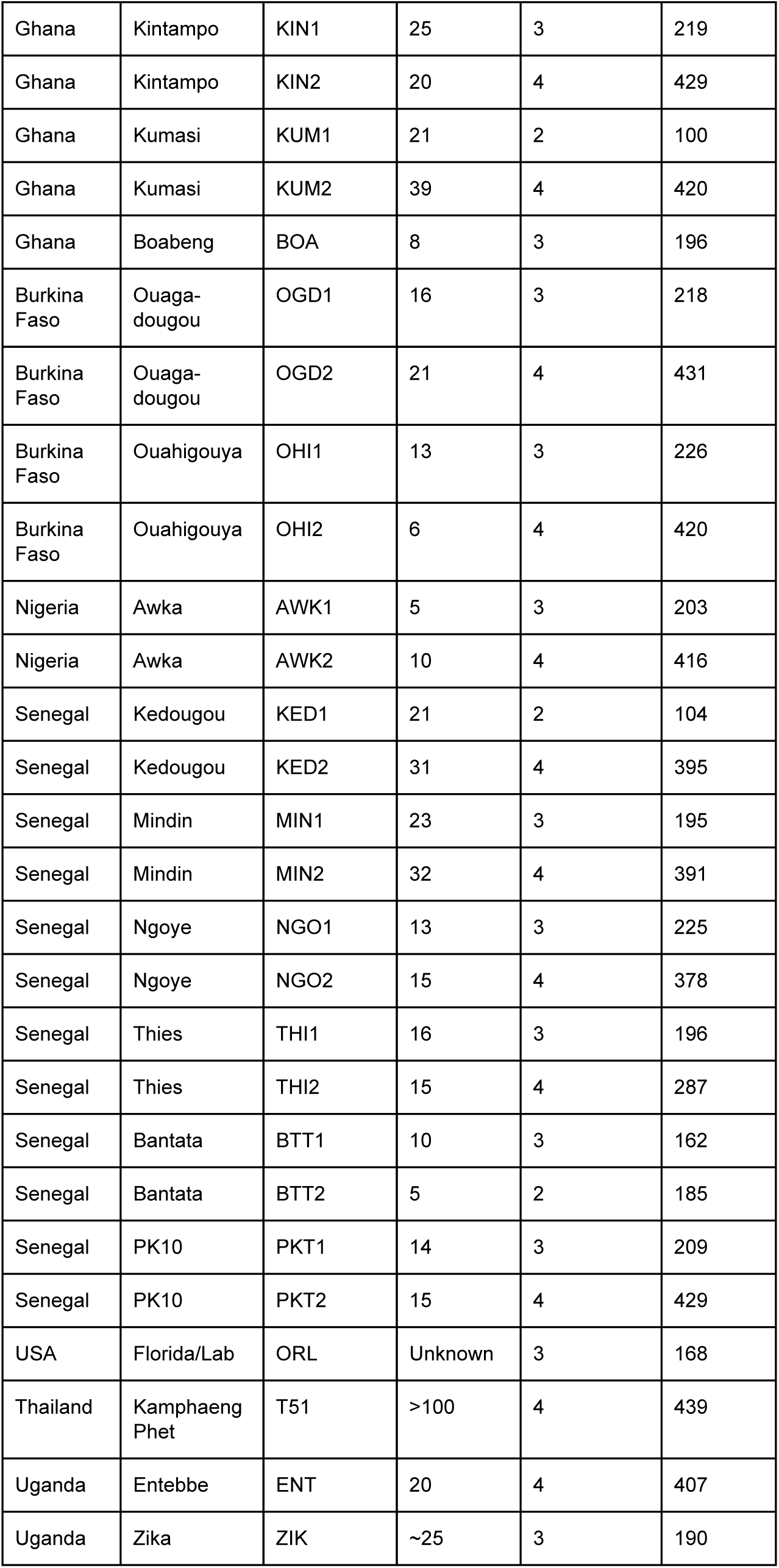
Details of laboratory colony establishment and behavioral testing.

